# Conformational Preference Classification of Integrin-Binding Ligands Using Free Energy Perturbation

**DOI:** 10.64898/2026.04.27.721214

**Authors:** Martin Vögele, Rezvan Shahoei, Loukas Petridis, Jing Li, Fu-Yang Lin, Lingle Wang, Timothy A. Springer, Jeremie Vendome

## Abstract

Integrins are crucial cell adhesion receptors and attractive therapeutic targets, but developing safe small-molecule inhibitors has been challenging, at least in part due to inadvertent partial agonism caused by stabilization of the integrin’s open, high-affinity state. To address this challenge, we present a computational approach using Absolute Binding Free Energy Perturbation (AB-FEP) calculations to predict whether a ligand will stabilize the open or closed integrin states, leveraging the difference between the ligand’s binding free energy to the respective end states. Despite challenges posed by Ca and Mg ions, metal-coordinating residues in the binding pocket, and the subtlety of structural differences between states, AB-FEP achieved excellent classification performance on a set of known opening and closing ligands, significantly outperforming docking scores and MM-GBSA results. We also showed a good correlation between AB-FEP binding free energy differences and experimental values. Furthermore, AB-FEP provided insights into intermediate integrin states and analysis of simulation trajectories confirmed the formation of a water-mediated hydrogen bond network with an ion in the binding pocket to be characteristic for closing ligands. This work demonstrates AB-FEP as a robust method for classifying integrin ligands by the conformation they stabilize and for understanding their functional mechanisms, offering valuable guidance for designing safe and conformationally selective integrin therapeutics.

**TOC Graphic:** 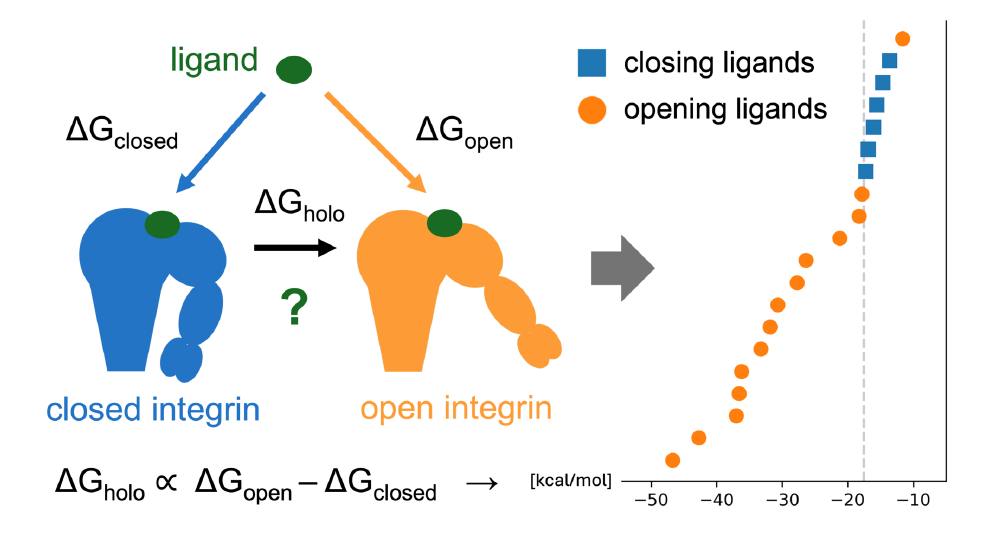

## INTRODUCTION

Integrins are cell adhesion receptors that bi-directionally signal across the plasma membrane and are involved in many physiological functions, including cell-to-cell communication, cell-to-extracellular matrix attachment, cell migration, and wound healing.^1–4^ Integrins have been shown to play an important role in many diseases like cancer,^5,6^ cardiovascular diseases^7,8^ and autoimmune disorders,^9^ which makes them attractive therapeutic targets.^10–12^ Currently approved small-molecule treatments include a topical medication for dry eye disease that targets the leukocyte integrin LFA-1 (αLβ2) and an intravenous acute treatment to prevent thrombosis that targets the fibrinogen receptor on platelets, integrin αIIbβ3. However, despite the emergence of new integrin targets in fields like fibrosis (αvβ6) and immuno-oncology (αvβ8),^13–18^ there are currently no approved oral small-molecule integrin inhibitors available.

After the success of parenteral small molecules approved to prevent thrombosis in patients undergoing per-cutaneous catheter procedures, many companies were optimistically developing oral integrin αIIbβ3 antagonists^19,20^; however, phase 3 studies showed the drugs made outcomes worse.^21–23^ Some evidence suggests inadvertent partial agonism: instead of purely blocking endogenous ligand binding, stabilization of integrins in their extended-open, high-affinity conformation may have triggered downstream signaling and adverse outcomes.^24–28^

Structural and biophysical studies on integrins have provided insights into the conformational changes underlying their activation (Figure 1).^29–31^ Integrin ectodomains have three overall conformational states, and on cell surfaces in the absence of ligand, about 99% of integrins are in the low affinity, bent-closed state, whereas in saturating ligand, most integrins are in an extended-open state. A series of crystal structures revealed the αIIbβ3 integrin in different intermediate conformational states, enumerated from S1 to S8 from the most closed to the most open state, bound to a soaked-in ligand that induced integrin opening.^32^ Biological ligands induce opening; however, a chemical principle to achieve a closing propensity was discovered for synthetic ligands that bind to the same site yet stabilize the closed state.^31^ Furthermore, clinical-stage integrin inhibitors that failed in trials were shown to stabilize the extended-open integrin state. Closure-stabilizing inhibitors shared a common chemical feature: a polar nitrogen atom positioned to hydrogen bond directly or indirectly with a gatekeeper water molecule (W1) located between the Mg^2+^ ion in the metal-ion-dependent adhesion site (MIDAS) and the side chain of a critical serine residue, Ser123 in the integrin subunit β3 (Figure 1C). This water acts as a gatekeeper for conformational change from the closed to the open conformation. During opening, the ligand-binding pocket tightens, and backbone movements at Ser123 toward the MIDAS Mg^2+^ ion bring the Ser123 sidechain oxygen into direct coordination with the Mg^2+^ ion and expel water W1. The hydrogen bond network between the polar nitrogen in closure-stabilizing compounds and the gatekeeper water are hypothesized to stabilize the receptor in its inactive, low-affinity state and replacing the polar nitrogen in compounds with carbon abolishes closure stabilization. Importantly, this chemical pattern was shown structurally and thermodynamically in αIIbβ3 and also thermodynamically with α4β1, therefore suggesting transferability across the integrin family.^31^ Thermodynamically it was shown that open-stabilizing compounds bind with higher affinity to the open than closed state, while closure-stabilizing compounds bind with higher affinity to the closed state. While hydrogen bonding to water 1 provides a strong predictor for the known ligands’ chemotypes for which structural data is available and it is clear whether such a hydrogen bond might be formed and with which part of the ligand, it might be more difficult to apply to new ligand chemotypes. Moreover, ligands with opening propensity show large variation in the affinity fold difference for the open and closed states, which the general chemical principle cannot capture.

**Figure 1.**
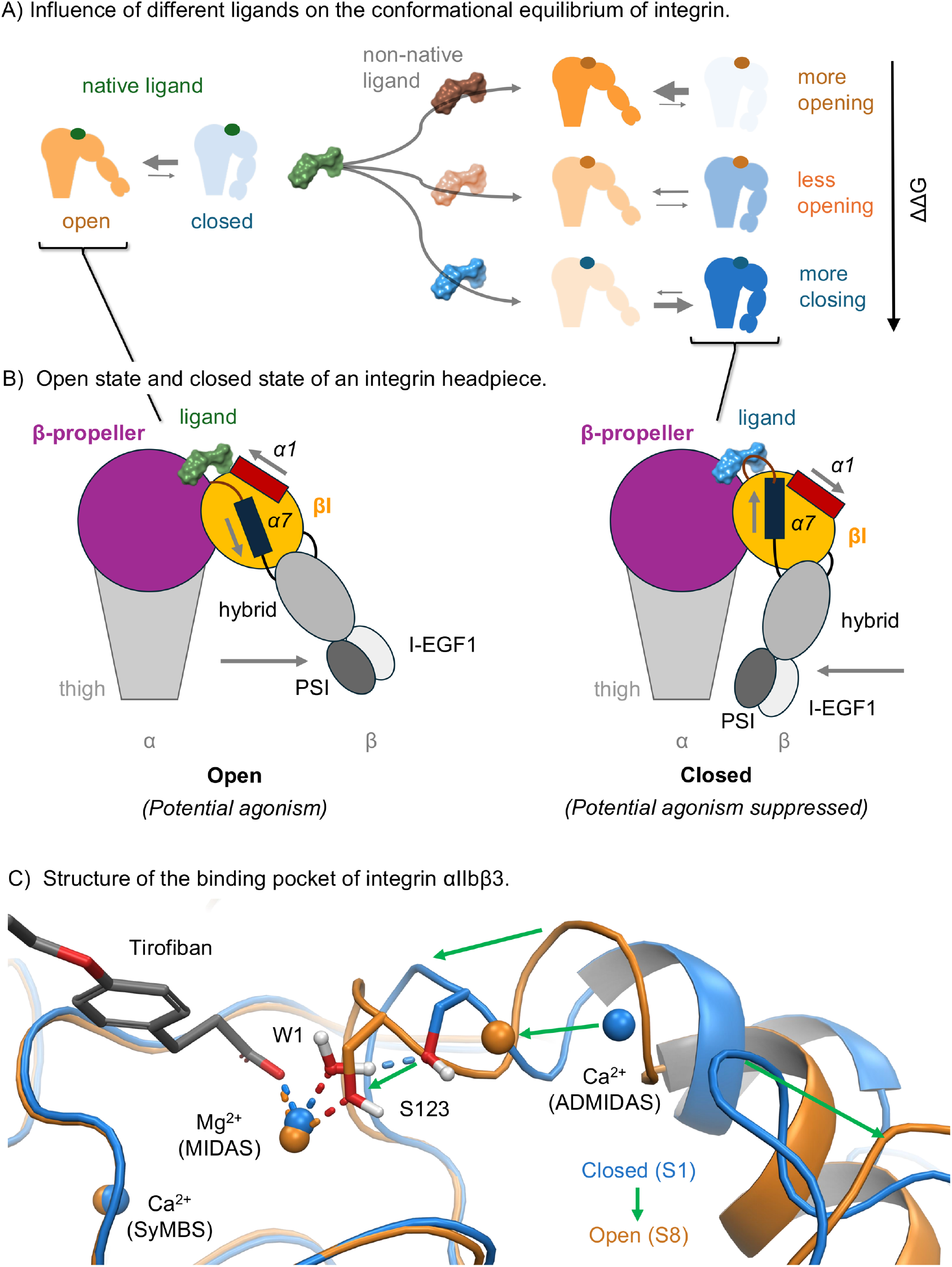
Integrins undergo conformational changes upon ligand binding; the equilibrium between the open and closed states, depends on the ligand’s propensity for these states. (A) Schematic of the influence of ligands with different opening or closing propensity on the conformational ensemble of an integrin headpiece fragment. (B) Schematic of the integrin headpiece. Our simulations include the β-propeller domain of the α subunit and the βI domain of the β subunit. (C) The βI domain ligand-binding pocket in the open (orange) and closed (blue) states. A part of the ligand tirofiban is shown in stick representation (carbons in gray) in the top left corner. Expulsion of the W1 water molecule that coordinates the MIDAS Mg^2+^ ion to Ser123 and is critical for the transition to the opening conformation. The green arrows highlight the movement for the ADMIDAS ion, the α1-helix, and the β6-α7 loop from the closed to the open state.

For these reasons, a computational approach that leverages these structural insights to predict whether a ligand has an opening or a closing propensity — and ideally the magnitude of this propensity — would help with prioritizing compounds in the development of safe and functionally selective integrin ligands. The propensity for one state or the other is theoretically linked to binding free energies, which are much easier to calculate than the reorganization of the entire protein. When a ligand binds to a receptor, it can shift the free energy difference between two conformational receptor states (Figure 1A/B) from ΔG_apo_ (opening free energy without the ligand bound) to ΔG_holo_ (opening free energy with the ligand bound). This ligand-induced shift, ΔΔG = ΔG_holo_ − ΔG_apo_ is equivalent to the difference between the binding free energies of the ligand to these two states.^33–36^ Specifically for integrins, the difference of binding free energy of a ligand to the open and closed conformations, ΔΔG = ΔG_open_ − ΔG_closed_, will determine whether it favors the open or closed state of the integrin (Supplementary Figure S1).^32^ Absolute binding free energy perturbation (AB-FEP),^37^ a computational method for predicting binding free energies, has been successfully used for a similar purpose: predicting the preference for a specific receptor conformation of ligands binding to GPCRs and a nuclear receptor.^36^ However, while FEP+ was applied with success to prioritize ligands for their affinity to the closed conformation in integrin αvbβ6,^15^ this approach has not been applied to predict the opening or closing propensity of integrin ligands so far. Integrins present a remarkable challenge for predicting a ligand’s opening or closing propensity through binding free energy perturbation due to two main issues: First, a general difficulty for molecular modeling arises from the presence of multiple strong charges in the binding pocket, specifically within the MIDAS and ADMIDAS regions, especially the non-covalently bound MIDAS and ADMIDAS metal ions themselves. Second, the structural differences in the binding pocket between the two states are quite subtle (Figure 1C).

In this work, we show that despite the many challenges of computationally predicting the opening or closing propensity of ligands, illustrated by the failures of other computational methods like docking and MM-GBSA (Molecular Mechanics - Generalized Born Surface Area), a more rigorous AB-FEP protocol achieved excellent performance in classifying compounds for their likely “opening” or “closing” propensity on integrin αIIbβ3, and can be used to identify early drug candidates with undesired partial agonist functional effect, and reduce failure in more advanced stages. Finally, we show how the investigation of FEP trajectories helped us understand the interaction patterns responsible for the opening or closing propensity of different ligands in integrins — a valuable source of insights for improved molecular designs.

## RESULTS

### Ligand Opening/Closing Classification Performance by AB-FEP

Predicting the opening or closing propensity of integrin-binding ligands using computational methods is a significant challenge, due to the very subtle structural differences in the binding pocket between the two conformational states (Figure 1C, RMSD ~ 1.1 Å, see Supplementary Table S2), as well as the presence of three metal ions in the proximity of the ligand-binding site. We evaluated our computational methods on a set of 16 integrin-binding ligands that are known to have an opening or closing propensity (10 and 6 ligands, respectively, see Supplementary Table S1). Their opening or closing propensity was determined experimentally by comparing binding affinities to the wild-type (WT) integrin αIIbβ3 and the mutant αIIbβ3 N305T which were used as proxies for the closed and open conformational states, respectively.^31^ Note that all the ligands actually bind to both conformational states, and that their opening or closing propensity reflect their relative binding affinity to the two states. For simplification, we will call ligands with opening and closing propensity, *opening* or *closing* ligands, respectively. For the assessment of the computational methods, we placed ourselves in a prospective context where the binding pose of the ligand in either of the two integrin conformations is unknown and used docking poses for all our calculations (see Methods section). We first employed the difference in docking scores to a closed-state structure and an open-state structure, a straightforward baseline approach for functional response prediction.^34^ This docking score difference, used as a classification metric, achieved an Area Under the Curve of the Receiver Operating Characteristic (AUC-ROC) of 0.50 (see Supplementary Figure S2 for detailed performance curves). This means that it did no better than a random classification and is thus not suited for robust classification in practical applications (Figure 2A). We also evaluated the performance of using binding free energy predictions from MM-GBSA calculations instead of docking scores, calculated from the poses obtained via the above-mentioned docking. Using binding affinity values from MM-GBSA yielded an AUC-ROC of 0.79 (Figure 2B), which is a significant improvement over docking scores. As MM-GBSA includes molecular mechanics and solvation energies (using an implicit solvent model) but not entropic contributions and explicit solvent, these results suggested that a more physically precise method, such as AB-FEP, could achieve even better results.

**Figure 2.**
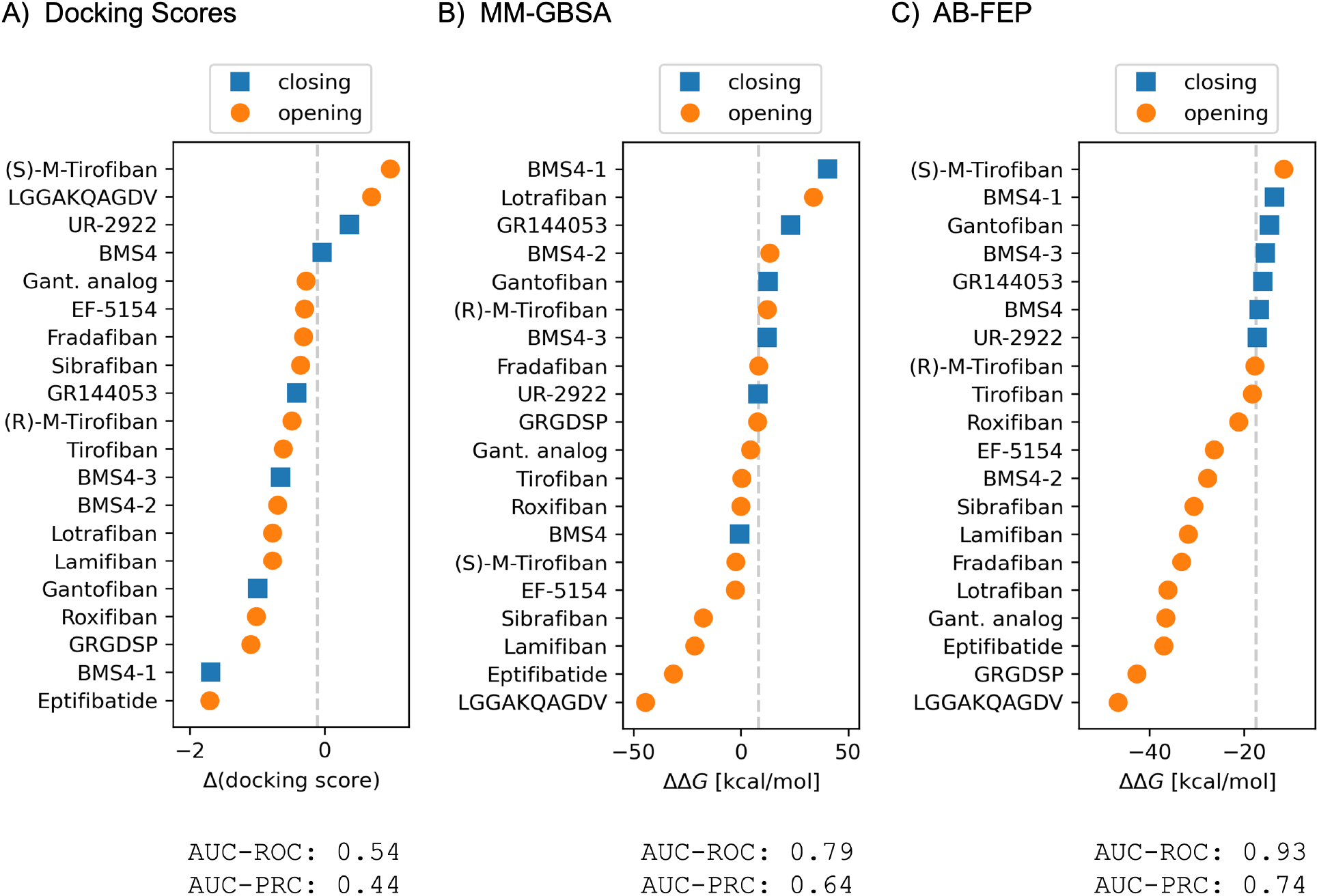
Absolute binding free energy perturbation with FEP+ successfully classifies integrin αIIbβ3 ligands as opening or closing. (A) Difference between the docking score to the open state and the docking score to the closed state, as obtained by Glide. (B) Binding free energy difference ΔΔG between the open and the closed state, obtained via MM-GBSA. (C) ΔΔG from AB-FEP with strong restraints to each state. For all three methods, State 8 (3ZE2-2) was used as the open state and State 1 (7TMZ-2) as the closed state, as these are the highest resolution structures for each state. In each panel, ligands are ordered by their increasing predicted likelihood to be opening and symbols and bars are colored by their actual opening property (blue: closing, orange: opening). The value for the optimal classification threshold is marked with a gray dashed line. The classification performance for each method is measured by the areas under the curve (AUC) of the receiver operating characteristic (ROC) and of the precision-recall curve (PRC).

The application of AB-FEP to ligands docked into both the open and closed conformations of the target protein yielded a binding free energy difference that demonstrated excellent performance as a classification score (Figure 2C), reaching an excellent AUC-ROC of 0.93 on the ligands evaluated in this study. Based only on the sign of ΔΔG, AB-FEP nominally would predict all ligands to bind more tightly to S8 (open), even the ones categorized experimentally as “closing.” Shifting the classification threshold from a theoretical default of zero to ΔΔG = −18 kcal/mol (Figure 2C) is necessary — and sufficient — to achieve the optimal classification. Instead of the difference ΔΔG, we similarly tried to use only one of the individual binding affinities ΔG_closed_ and ΔG_open_ as a metric along which to separate opening and closing ligands. For ΔG_closed_, this yielded an AUC-ROC of 0.76 and, analogously, the AUC-ROC for classifying a ligand as opening using only ΔG_open_ was 0.73 (see also Supplementary Figure S10 E/F). This means that, while binding to a single state alone can yield some insights, the difference in binding free energy is needed for optimal classification. A particularly encouraging result is that ΔΔG from AB-FEP correctly predicted the opening or closing propensity within the small congeneric sets present in our dataset — namely between gantofiban (closing) and the gantofiban analog (opening), and between BMS4 and its three analogues. Such ability to predict structure-activity relationships within congeneric series, where the chemical changes are typically subtle, is particularly relevant in the context of drug design, for the optimization of a given compound series. Ultimately, the AB-FEP protocol we proposed proves to be remarkably effective and reliable for the classification of integrin-binding ligands.

A crucial aspect of the AB-FEP protocol are strong harmonic restraints on the protein backbone atoms during the FEP simulations in the two respective conformational states, motivated by the subtle structural distinctions between the two protein states and intended to avoid too much overlap in the conformational space sampled during the FEP simulations. To examine the impact of different methodological choices, we explored different AB-FEP protocols. For these and all subsequent simulations, we used experimental poses instead of docking poses, as they are available for all the studied ligands except (R)- and (S)-M-tirofiban in at least one of the eight known integrin conformational states. This way, we avoided the need to simulate and select the best pose from multiple docked options. When using our standard protocol, calculations started from the experimental poses exhibited similar accuracy (AUC-ROC: 0.92; Supplementary Figure S3) as the docked poses. When we used wider restraints of the protein backbone (flat bottom of 1 Å, see Methods), the AUC-ROC was reduced to 0.78 (Supplementary Figure S4), which emphasizes the importance of a very narrow conformational sampling around each state for the case of integrins. Another important aspect of the AB-FEP protocol proposed is the use of metal custom charges for the OPLS4 force field including zero-order bonds to the metal ions (see Methods) due to the presence of strongly charged ions within the binding pocket.^38–40^ Without these metal custom charges, we only reached an AUC-ROC of 0.75 (Supplementary Figure S5). These results collectively validate our methodological decisions to use custom partial charges and strong protein backbone restraints.

### Correlation between Predicted and Experimental Binding Affinity Difference

Beyond binary classification, we found that the binding preference predicted by our AB-FEP protocol and quantified by the magnitude of the predicted free-energy difference to the two integrin states, correlated with the experimentally derived values, namely the difference between the experimental binding affinities to the (closed proxy) wild-type (WT) integrin αIIbβ3 and the (open proxy) mutant αIIbβ3 N305T.^31^ While a scaling factor is necessary to align the absolute magnitudes, reflecting that the absolute free energy values and their corresponding differences obtained from our AB-FEP calculations do not directly correspond to the experimental values, the correlation with experimental values is much higher than for docking scores and MM-GBSA: R^2^ = 0.42 compared to 0.14 and 0.03, respectively (Figure 3A–C). Note that we excluded the linear peptides from calculating the correlation because they were simulated with a longer simulation protocol. Additionally, the linear peptides might need a different offset or scaling factor but, with only two peptide data points, we cannot calculate a separate offset and scaling factor — or even determine whether it is necessary. A general limitation of this performance evaluation is that the experimental ΔΔG values correspond to measurements on ensembles that approximate but are not pure closed and open ensembles. Despite the significant deviations in the absolute binding free energy values, it is particularly noteworthy that the (scaled and shifted) *difference* in free energy between the two conformational states allows for a robust and reliable assessment of the ligand’s preference for one of the two major conformational states of integrin.

**Figure 3.**
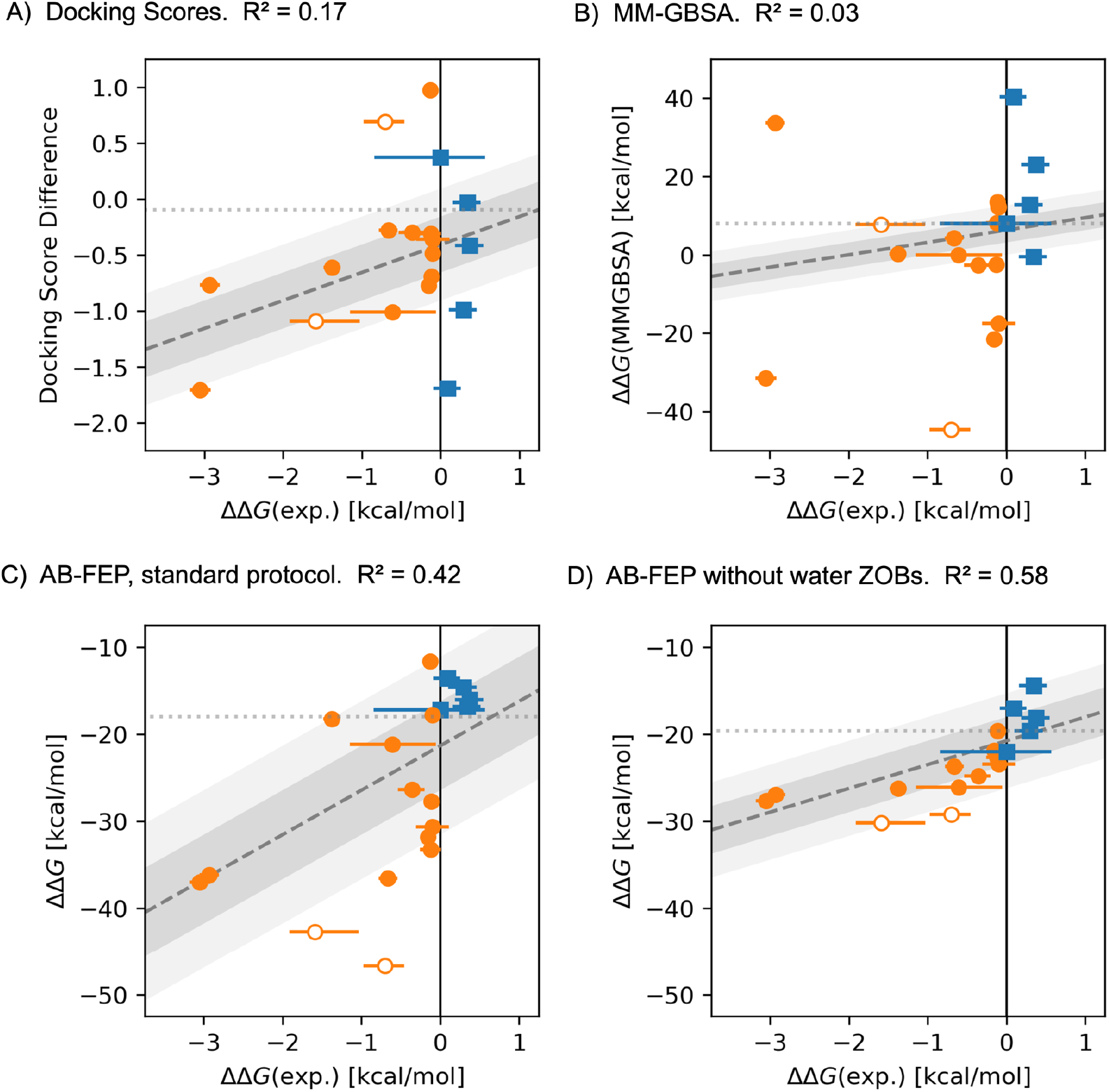
Correlation of classification scores from various methods to the binding free energy difference between the open and closed state obtained from experiments. All computational methods used the structure from 7TMZ for the closed state and from 3ZE2 for the open state. Experimental binding assays used the WT of αIIbβ3 for the closed state and the N305T mutant for the open state. The long-dashed line shows the least-squares fit with gray shaded error bands corresponding to 1 kcal/mol and 2 kcal/mol in the experimental ΔΔG. The short-dashed horizontal line shows the optimal threshold for classification. The two linear peptides in the dataset are depicted as empty circles and excluded from the calculation of the R^2^ reported.

A possible limitation to obtaining an even better correlation could be the lack of flexibility for the waters coordinating the MIDAS ion to adjust to different ligands, due to the usage of zero-order bonds (ZOBs) between these water molecules and the metal ions in our protocol, thus restraining both distance and angle of the water molecules relative to the ion and each other. This hypothesis is supported by a control test using OPLS4 with metal custom charges (OPLS4-mcc) without zero-order bonds between metal ions and water molecules (while retaining protein-metal ZOBs). This modification further improved the correlation between the predicted and experimental ΔΔG values to R^2^ = 0.58 (Figure 3D, Supplementary Figure S6), as well as the correlation of the predicted affinity to each individual state with the corresponding experimental state (Supplementary Figure S11).

We should emphasize, however, that this is an *ad-hoc* modification that we do not recommend as a general solution with OPLS4-mcc. This force field was parametrized accounting for the presence of ZOBs, which were found to be necessary in other metal-coordinating systems to maintain a stable coordination geometry. In an additional set of simulations, removing *all* ZOBs led to similar results but for the decapeptide the active-state simulation was too unstable to complete (Supplementary Figure S7). Taken together, these results support our interpretation that the lack of flexibility of the water molecules due to ZOBs limits the accuracy of the correlation between the predicted and experimental ΔΔG values.

### Intermediate Integrin States

To further investigate the structural determinants of the signal captured by our AB-FEP protocol and allowing to distinguish between opening and closing ligands, we used an intermediate-state integrin conformation (PDB 7UDG, molecule 1, see Figure 4A).^32^ This intermediate state is the result of the soaking process required to obtain the crystal structures. The integrin was crystallized in an inactive state in which its global conformation remained trapped. The soaked-in agonist could only locally adapt the binding pocket because the crystal lattice prevented global conformational changes. Thus, such a state allowed us to study the effect of local conformational signatures of agonists in the binding pocket. The structure used here was classified as S3 via the geometric classification — in particular the position of the ADMIDAS — that classifies S1 (7TMZ-1) and S8 (32EZ-2) as the nearest to the closed and open states, respectively.^32^ Following this geometric state definition, the intermediate-state conformation from 7UDG-1 (S3) is expected to be more similar to S1 than to S8. However, while the overall structure of integrin in this structure (S3) is indeed mostly closed (global RMSD of 0.21 Å and 1.53 Å to S1 and S8 respectively, Table S2), the direct vicinity of the ligands in the binding pocket resembles more that of an open state (local RMSD of 0.74 Å and 0.46 Å to S1 and S8 respectively, Table S2), especially for the loop including Ser123 (Figure 4A). This corresponds to a step change in the coordination of the MIDAS ion from water 1 (S1 and S2) to Ser123 (S3 to S8), which is why we chose S3 as the most interesting intermediate state to study the effect of local vs global rearrangements. We then used our AB-FEP protocol to predict the opening or closing propensities of our ligand dataset, but this time substituting the S3 state for either the closed state (S1) or the open state (S8). The performance decreased drastically when we attempted classification using S3 instead of the fully closed-state (S1) structure while keeping the fully open-state structure (32EZ-2) (Figure 4B, detailed results in Supplementary Figure S8). Remarkably, when substituting S3 for the fully open state (S8) structure and keeping S1 for the closed state, we obtained similar performance as with the fully open-state structure (Figure 4C, detailed results in Supplementary Figure S9). Furthermore, it appears that the individual affinities predicted for the dataset ligands to the S3 intermediate-state conformation, are much closer to affinities predicted on the S8 state than the S1 state (Figure 4D). They also correlate negatively with wild-type affinities and slightly positively, yet still much worse than S8, with mutant affinities (Supplementary Figure S12). This strongly suggests that the details of the integrin conformation in the direct vicinity of the ligand are the most important for the classification performance in our AB-FEP protocol, which is consistent with the restraints used in it.

**Figure 4.**
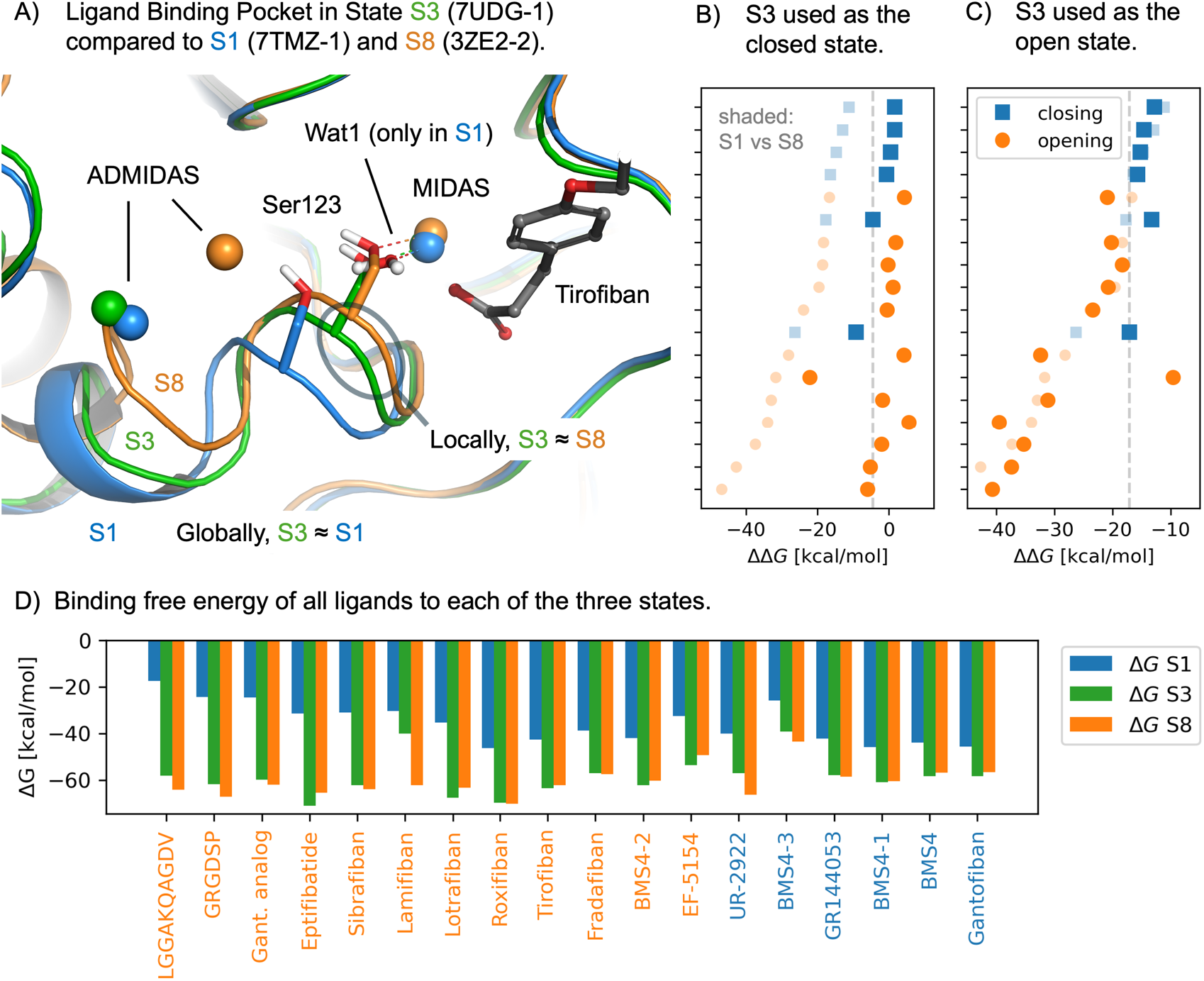
Tests of AB-FEP on state S3, which is locally similar to the open state in the ligand-binding pocket and globally similar to the closed state. (A) Overlay of the three states S1, S3, and S8 (blue, green, and orange carbons and metal ions, respectively), with tirofiban (grey carbons) for orientation (sulfonamide-linked moiety omitted for clarity). The Mg^2+^ ion at the MIDAS and the Ca^2+^ ion at the ADMIDAS are shown as spheres. Similarities between the structures are quantified in Supplementary Table S2. (B) Classification of integrin binders if we use S3 as the closed state and S8 as the open state compared to using S1 and S8 (shaded small symbols). Ligands are ordered by their ΔΔG obtained from S1 and S8. This combination of states provides no predictive value. (C) Classification of integrin binders if we use S3 as the open state and S1 as the closed state compared to using S1 and S8 (shaded small symbols). Ligands are ordered by their ΔΔG obtained from S1 and S8, as in panel B. S3 yields results comparable to those obtained when S8 is used as the open state, suggesting that its ligand interaction behavior is similar to an open state. (D) Binding free energy of all ligands to each of the three states. Opening ligands are labeled orange and closing ligands are labeled blue.

### Water Hydrogen Bond Patterns of Opening and Closing Ligands

The analysis of the simulation trajectories from the AB-FEP calculations confirmed the presence of a water-mediated hydrogen bond pattern with the MIDAS ion to be a hallmark of closing ligands, in the closed state specifically. Lin et al. observed that X-ray structures of closing inhibitors bound to the closed integrin state consistently showed a water-mediated hydrogen bond network connecting a nitrogen atom within the ligand to Ser123.^31^ The highly successful categorization of the ligands with the AB-FEP approach reported here, which rigorously accounts for the affinity of the ligands for both end states, as well as some level of dynamics in the structures, was an opportunity to better understand this pattern. Indeed, in our AB-FEP simulations a hydrogen bond network that includes water 1 connects the MIDAS ion with a nitrogen in the ligand in the closed state S1 (Figure 5, Table S3) for 6 out of 6 closing ligands, independent of whether the protonated or deprotonated state of the nitrogen dominates for a particular ligand. The protonated form of the nitrogen makes it possible for only one water molecule to bridge the gap between the ligand and Ser123. In the deprotonated state, two water molecules are needed to bridge the gap. For BMS4, the protonated state dominates according to population analysis with EpikX (90% vs 10%), however, the network with two water molecules bridging deprotonated BMS4 to Ser123 (Figure 5C) agrees better with its experimental structure (PDB: 7TMZ) than the single water molecule in the simulation of protonated BMS4 (Figure 5D). In contrast to closing ligands, most opening ligands in our simulations lack a hydrogen bond network to water 1 and the MIDAS ion in the closed state S1 (Figure 5, Table S3). With our simulations, it is now possible to also investigate the behavior of all the ligands in the open state. We found that none of the closing ligands exhibited this hydrogen bond network in the open state, which explains why they prefer the closed state in which their carboxyl-proximal nitrogen can form such a network. Of the opening ligands, three did show a hydrogen bond network in the open-state simulations (Supplementary Table S3). This could contribute to those particular ligands having an opening propensity — but it does not appear to be a general pattern. Overall, the presence of a hydrogen bond connection between ligand, water 1, and Ser123 in the closed state is a good indicator of whether a ligand is closing or opening.

**Figure 5.**
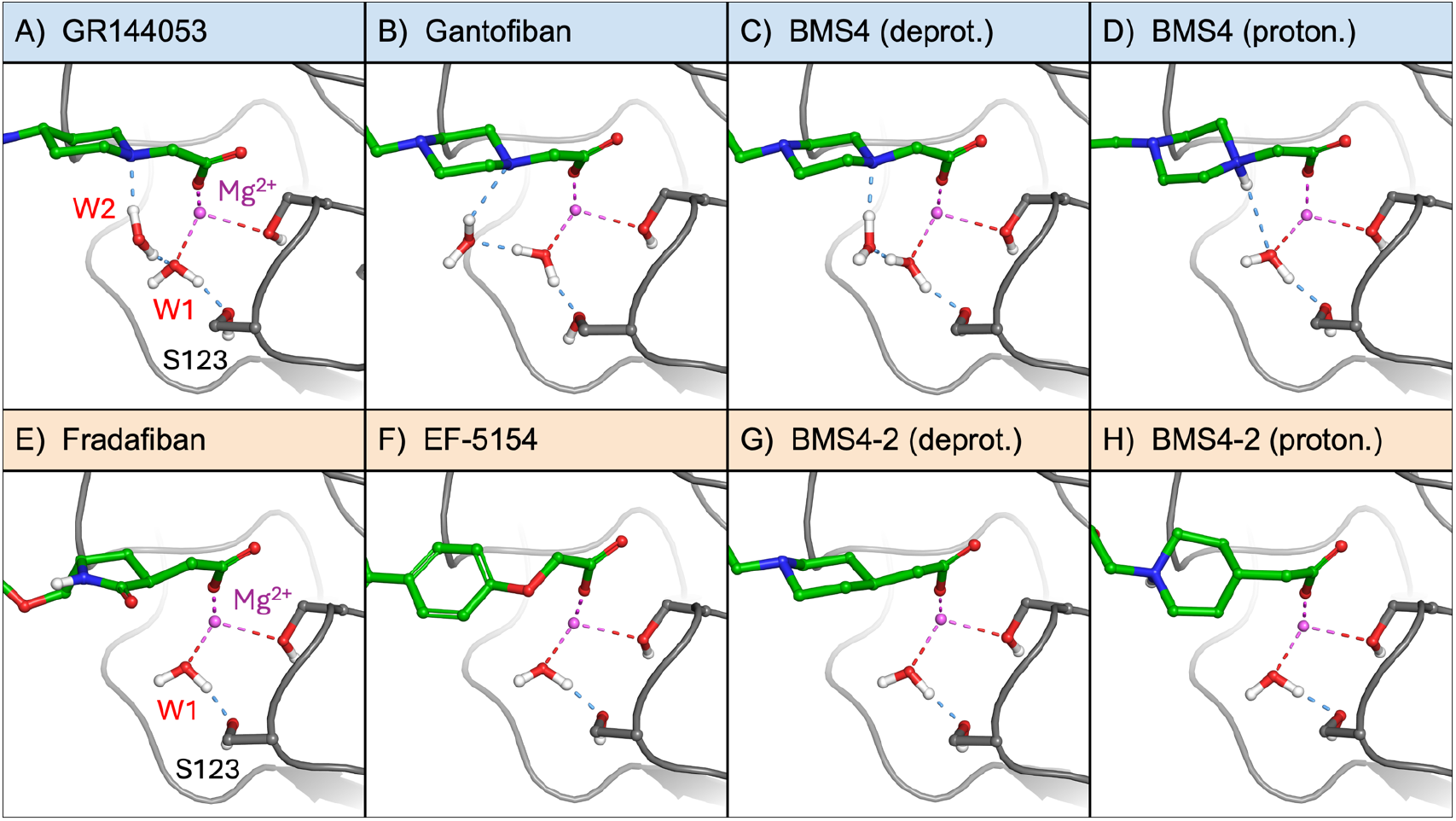
The hydrogen bond pattern in the closed state S1 during the AB-FEP simulations differs for closing ligands (A–D) and opening ligands (E–H). The hydrogen bond pattern for BMS4 involves one water molecule with protonated nitrogen and two waters with deprotonated nitrogen. Representative structures from AB-FEP of the S1 state are shown, with hydrogen bonds in light blue, the MIDAS ion in magenta, and the protein in gray. Results for all the ligands are shown in the Supporting Information Table S3.

## DISCUSSION

The excellent classification performance we achieved using AB-FEP demonstrates that the difference between binding affinities to open and closed states is a robust predictor of whether an integrin binder has an opening or closing propensity. The thermodynamic principle that functional response of a ligand can be linked to its binding affinities to the functionally relevant states – as shown theoretically and for GPCRs and nuclear receptors – also works for the more challenging integrin. Note that, unlike for the previous AB-FEP study, the experimental ground truth for the ligands in the integrin dataset used here is biophysical, determined directly from binding affinity to wild-type or mutant receptors, rather than from a functional assay.^31^ A particular difficulty here, in contrast to the other systems for which such a workflow has been tested previously, is that integrins have three divalent cations in the binding pocket, giving an outsized role to electrostatic interactions. Also, the structural differences in the binding pocket between the two states are small. The restraints needed to avoid overlapping conformational ensembles had to be narrower than in earlier work on GPCRs and NRs (the latter of which did not need restraints at all).^36^

Importantly, the classification accuracy obtained with AB-FEP was similarly good when using experimental complex structures, and when using a docking protocol more representative of active drug discovery efforts where structural information might be limited. In cases where experimental complex structures are available for a ligand series of interest, usage of these structures is still recommended as certain ligands can induce specific binding pocket rearrangements relative to available integrin structures. For example, RUC-4 even displaces the metal ion at the MIDAS.^41^ In one case, UR-2922, we obtained a better classification with docked poses than with the experimental pose, even though the docked pose differed significantly because Glide placed the flexible arm with the nitrophenyl group in the opposite direction (Figure S16). Since this occurs in both states, it is likely an example of fortuitous cancellation of errors. This kind of effect can make predicting ΔΔG more robust than individual ΔG and even than the underlying poses. As the only classification outlier, the propensity for the (S)-M-Tirofiban enantiomer is mispredicted by our protocol using the docked pose — while its (R)- enantiomer is well in line with the other results. Unfortunately, we cannot assess whether this misclassification is due to an inaccurate docked pose, as this ligand does not have an experimental crystal structure available for comparison. Yet, in cases where no complex structure was available, the present study demonstrated that using a docking protocol with AB-FEP can be a very valuable alternative.

AB-FEP exhibited superior performance to MM-GBSA in a realistic scenario using docking poses. The success of AB-FEP was due to two main reasons: First, AB-FEP explicitly models the solvent, capturing the dynamic nature and precise energetic contributions of hydrogen bonds, unlike MM-GBSA’s implicit solvation model which approximates these effects. Second, AB-FEP natively incorporates entropic effects by thoroughly sampling conformational space and directly calculating the free energy difference, whereas MM-GBSA relies on approximations for entropy that may not fully capture the binding process. Thus, beyond successfully validating the thermodynamic principle that underlies the previous successful prediction of functional response on other systems, we demonstrated that AB-FEP is an effective method for applying this principle to opening and closing propensity on integrin, as it accounts for many important effects neglected by other methods.

Predicting a ligand’s preference for a specific functional state is more robust than predicting the absolute binding free energies. The individual binding free energies predicted *via* AB-FEP are much larger than the experimentally measured values. This is due to several inherent challenges in calculating precise free energy values. One of the primary reasons for discrepancies in free energy calculations stems from the force field parameterization. The OPLS4 force field with metal custom charges was parameterized without the ligand present during this process. Even if it were technically feasible to parameterize OPLS4-mcc in the presence of the ligand, the partial charges on binding pocket residues would undergo significant changes between the bound and unbound states within a single FEP calculation. Such a dynamic adaptation of the force field is technically complex and is not currently implemented in FEP+. Furthermore, strong electrostatic interactions, which are often critical in ligand-protein binding, are particularly susceptible to even minor inaccuracies in the underlying force field and simulation parameters. These interactions are highly sensitive to the precise atomic charges and dielectric properties, making their accurate representation a significant challenge. In addition to these challenges stemming from the force field parametrization, the conformational sampling was also likely biased by the restraints that were found to be necessary for optimal classification. In particular, the restraints on the water molecules coordinating the ions (ZOBs) might have amplified the inaccuracies for the electrostatic interactions. Given these considerable difficulties and technical limitations, it is remarkable that the opening or closing propensity of the ligands can be predicted with such a high degree of accuracy. A likely explanation for this unexpected success is that many of the inherent errors in the individual free energy calculations tend to cancel each other out when focusing on relative preferences rather than absolute values. This shows that AB-FEP can yield meaningful insights into relative binding preferences, even in scenarios where precise absolute predictions remain challenging.

While in general a theoretical classification threshold of ΔΔG = 0 is expected theoretically the systematic shift in the boundary necessitates an empirical determination for each pair of template structures. This threshold shift is usually the result of multiple factors that can result from computational modeling or the underlying experimental structures. In the case of integrin, the main cause is likely the strongly polarized interactions coordinating the doubly charged ions. As discussed above, force field representations for these interactions are challenging to parameterize, making it difficult to achieve a consistent treatment across different conformational states. In particular, the transition between a protein-ion bond and a protein-water bond from one structure to another introduces unphysical energy changes that are not perfectly balanced by partial charge adaptations. We think that this effect is particularly critical at the transition from MIDAS coordination by water 1 to coordination by Ser123 because the threshold changes very little from classification using States S1 and S8 (Figure 2C) to using S1 and S3 (Figure 4C) but a lot in the failed attempt to use S3 and S8 (Figure 4B). Prior investigations involving GPCRs identified instances where an empirical shift in the classification threshold was caused by structural artifacts, such as inadequately resolved loops or non-physical deformations from crystal packing interactions that caused an underestimation of binding affinity for one of the two states.^36^ We cannot exclude that such artifacts play an additional role here. Since it is difficult to disentangle the individual contributions of each possible factor, the adapted threshold accounts for all of them together. In practical applications, the classification threshold should be gauged for a specific pair of template structures using a small set of known ligands before scaling up the prediction to large prospective datasets. The same shifted threshold as determined on a set of two integrin structures would then be applicable to different ligand chemotypes. However, it would not translate to different integrin states and different integrin types. For a new integrin structure, the partial charges and zero-order bonds need to be re-parameterized which can shift the artifacts introduced by these in a different way in each state. Thus, the classification threshold must be determined for each pair of structures anew.

While the correlation with the experimental ΔΔG is not as strong as desired, it is sufficient in the affinity range most important for classification. Particularly within the opening ligands, the correlation across the small molecules in this dataset is weak. A potential source of artifacts is the use of ZOBs, which may be overly restrictive. ZOBs not only prevent the adaptation of equilibrium distances and angles in the water network to the presence or absence of a ligand but also reduce their fluctuations and thus alter the entropic component of the binding free energy. The extent of these effects and the resulting artifacts can vary by ligand. Indeed, omitting the ZOBs involving water improved the correlation (Figure 3D). As an extreme example for a structural rearrangement prevented by ZOBs, we observed in one simulation without water ZOBs that the reordering of water molecules even resulted in a flip of the sidechain of Ser123 (Figure S15). While these results may tempt us to omit ZOBs, it should be cautioned that this could have unforeseen effects on other relevant properties since, with OPLS4, ZOBs are necessary to maintain a stable geometry around metal ions. We thus do not recommend removing ZOBs as a general solution and chose to retain them in our standard protocol which, after all, yielded excellent classification performance. Future iterations of the OPLS force field could address the conflict of aims between conformational flexibility and accurate partial charges by modeling ion polarization effects on their surroundings without the necessity of ZOBs. Such improvements might also resolve the remaining deviations from optimal correlation if they are caused by ZOBs between ions and protein and improve our ability to quantitatively predict the opening propensity of different ligands In particular, this can help answer the open questions of why opening ligands vary so much in the degree of their opening propensity and whether biological integrin ligands have evolved to exhibit a high opening propensity. On the experimental side, we should keep in mind that the experimental ΔΔG does not exactly correspond to ΔG_open_−ΔG_closed_, as we used the affinity for the N305T glycan wedge mutant to approximate the intrinsic affinity for the open state, although this mutant is not fully open.^31^ The actual ΔΔG can be considered to be more negative for the open compounds, and more positive for the closure compounds. Accounting for this effect quantitatively is not possible but we would expect it to increase the correlation R^2^. Further deviations could be caused by differences in kinetics that might cause the extent of integrin opening to vary among ligands. While it appears to remain challenging to distinguish weakly opening from strongly opening ligands, our workflow is well suited for identifying closing ligands, which for the purpose of drug discovery is of a higher priority.

The signal captured by our AB-FEP protocol depends much more on the local conformational difference between states in the proximity of the ligand in the binding pocket than on the global conformational differences and overall shape of the receptor, as we show by probing the intermediate integrin state S3. Despite its structural resemblance to the closed-state structure (S1), the binding free energies predicted using this intermediate state align more closely with those predicted in the fully open integrin state S8, albeit not perfectly. This result highlights the importance of the binding pocket’s local conformation, especially the loop close to the ligand and the MIDAS ion over the receptor’s overall state. It also exemplifies a general warning for binding free energy calculations: a receptor might appear to be in one functional state when examined globally, but small conformational details, especially in its binding pocket, could make it more aligned with another functional state for free energy calculation purposes. Such intermediate structures might be stabilized by specific ligands with particular state preference. The emergence of the intermediate states in these structures can be explained by the soaking crystallization process used, where integrin is initially in a closed state which is maintained by the inherent restraints of crystal packing, even when ligands are introduced. In this scenario, ligands can only adjust their immediate surroundings within the binding pocket, even if they would otherwise induce an open integrin state in solution. This is corroborated by the observation that intermediate states like S3 can only be observed in crystal structures of opening ligands.^31,32^ While intermediate states like the S3 structure probed here can yield interesting mechanistic insights, the best combination of states to use for classification purposes is still the most straightforward one: a pair of the most extreme states, resolved with matching ligands.

While theoretically two calculations are needed to classify ligands as opening or closing, practical use cases do not require all ligands to be tested this extensively. To streamline the process, binding to the preferred state should be investigated first. For example, when designing closing inhibitors, candidates should initially be screened for good binding to the closed state. Beyond ensuring an overall strong affinity, this initial screening also helps enrich for ligands with a closing propensity, as evidenced by an AUC-ROC of 0.81 for classifying a ligand as closing using only the AB-FEP affinity to the closed state (at least for the subset studied here). For ligands sharing a common scaffold, this step can also be performed using the computationally more efficient relative binding FEP.^42^ Subsequently, only the most promising candidates need to be tested additionally for binding to the open state to calculate the predicted propensity.

Beyond predicting the opening or closing propensity of a ligand, free energy perturbation simulations can also provide insights into the underlying mechanism. The analysis of the trajectories allowed us to confirm the hypothesized role of a water-mediated hydrogen-bond network connecting Ser123 to a nitrogen atom in the closing ligands among the investigated set of integrin binders. Specifically, we could confirm the consistent observation of such a water-mediated hydrogen bond network during the simulations in the closed state for all closing ligands, but not for opening ligands. And importantly, this hydrogen bond network is mostly absent for all ligands in the simulations in the open state. The importance of hydrogen bonds to and among water molecules makes explicit solvent water indispensable in accurately capturing the system’s dynamics. The presence of explicit water molecules is a primary reason why AB-FEP outperforms other computational approaches that omit explicit solvent. The nitrogen atom capable of forming the characteristic hydrogen-bond pattern seen in the closing ligands has no equivalent in the tested peptides. Even though they have nitrogen atoms, those are positioned further away from the anchoring carboxylate group, which likely explains why they are consistently opening. Ultimately, the application of AB-FEP extends far beyond mere predictive capabilities by providing mechanistic insights based upon which to conceive and design novel molecules with improved efficacy.

## CONCLUSION

We present a workflow based on AB-FEP to predict whether an integrin-binding ligand is opening or closing. Integrin is a challenging system for this task, as demonstrated by the failure of our baseline approach of simply comparing docking scores to open and closed states. We found that MM-GBSA performed better than docking scores but did not reach satisfactory accuracy. Even in AB-FEP, the absolute values were off, and the classification threshold was shifted. Despite these challenges, we succeeded in using AB-FEP for classifying integrin ligands, showing that opening/closing propensity prediction is robust. By varying partial charges and zero-order bonds, we explained the factors that enhance and limit performance of our approach. We also confirmed the structural hypothesis that closing inhibitors favor a water molecule that bridges a serine residue and a metal ion in the MIDAS and gained insights into a change in the dominant protonation state. In summary, we demonstrated a workable solution for predicting whether integrin binders are opening or closing and can provide valuable insights into the corresponding mechanism.

## METHODS

### Ligands and Experimental Reference Data

We investigated a set of ligands for which the opening or closing propensity were determined experimentally via their binding affinities for αIIbβ3 WT and αIIbβ3 N305T — a mutant proxy for the open state (Supplementary Figure S14 and Supplementary Table S1).^31^ For classification studies, we included the closing small molecule BMS4-3 to reduce the imbalance in the number of closing and opening ligands, though it cannot be included in correlation analyses due to its unmeasurably weak binding affinity. We did not include the neutral (neither opening nor closing) compound RUC-4 because its mechanism of action involves displacing the metal ion at the MIDAS,^41^ making it unique among the ligands with available experimental data. The constructs used in our workflow, have a metal ion fixed that cannot change during the simulation (see System Setup).

### System Setup

Eight distinct conformations for αIIbβ3 integrin headpiece starting from the closed state (S1) to the open state (S8) have been previously reported.^32^ Aside from one closed and two open states, we also built a model for an intermediate state (S3). The following PDB structures were used to build these models: molecule 1 of 7TMZ (7TMZ-1) reported in state 1, molecule 1 of 7UDG (7UDG-1) reported in state 3, molecule 2 of 3ZE2 (3ZE2-2) reported in state 8, and 2VDM also reported in state 8. In all cases, the Fab heavy and light chains were removed. Chain A in 7TMZ-1 and 7UDG-1 and chain C in 3ZE2-2 were aligned to chain A in 2VDM. To ensure smaller system sizes for calculation efficiency, the residues in chain B/D outside the range of 108–343 were truncated.

The metal ions in the binding pocket warranted special attention. Among the above four crystal structures, 7TMZ-1, 7UDG-1, and 2VDM were solved with Mg^2+^/Ca^2+^. There is one Mg^2+^ ion at the metal-ion dependent adhesion site (MIDAS), one Ca^2+^ ion at the adjacent to MIDAS (ADMIDAS) and one Ca^2+^ at the synergistic metal binding site (SyMBS). 3ZE2-2, on the other hand, was solved with three Mn^2+^ ions, one Mn^2+^ ion at each site. The Mn^2+^ ions in 3ZE2-2 were manually mutated to Mg^2+^ for the MIDAS ion and Ca^2+^ for the ADMIDAS and SyMBS ions. The ADMIDAS Ca^2+^ is missing in the 7UDG-1 structure. We manually built it using the 7TMZ-1 ADMIDAS Ca^2+^ as the template. Zero-order bonds were built such that the coordination geometry for the ADMIDAS Ca^2+^ in the modeled systems is consistent with previously reported findings, i.e., the coordination number of 7 (pentagonal bipyramidal geometry) in the closed state and coordination number of 6 (octahedral geometry) in the open state.^43^ The coordination number for the built-in ADMIDAS Ca^2+^ in 7UDG-1 (S3) was assumed to be 7, similar to the closed state (S1). In 2VDM, the ADMIDAS Ca^2+^ in the crystal structure coordinates with a glycerol, a crystal artifact. The glycerol was replaced with two water molecules as coordinating partners for the ADMIDAS Ca^2+^.

All structures were modeled and prepared in Maestro.^10^ The Protein Preparation Workflow^44^ was used to fill in missing side chains, to cap all termini, and to generate the likely protonation and tautomeric states with Epik at pH 7.4 ± 2.0. Using PROPKA, all protonation states at pH 7.4 were determined such that the hydrogen bond network was optimized.

### Force Field Parameterization

Due to the high charges involved in the metal coordination within the binding pocket and to better represent charge transfer effects between the metal and coordinating groups, we made adjustments to the standard OPLS4 force field.^38–40^ In all the ligand-receptor systems in this study, the binding pocket is highly charged and the MIDAS Mg^2+^ coordinates with the ligand’s carboxylate group. At the time of running the calculations, OPLS5 with metal support was not yet available and, therefore, we used an early development prototype of OPLS5 with metal support, *i*.*e*., OPLS4 with QM fit partial charges for the metal coordination complex. This early development prototype will hereafter be referred to as OPLS4-mcc for metal custom charges. It is worth noting that realistic characterization of metal coordination using classical force fields is challenging. In OPLS4-mcc, the issue is handled by “zero-order bonds” (ZOBs)^45^ between the metal and the coordinating partners where the input coordination geometry is maintained via bond stretch/angle constraints. OPLS4-mcc was used in all MM-GBSA and AB-FEP calculations except for one AB-FEP run where OPLS4 was used for comparison.

To determine the metal custom charges for OPLS4-mcc, first a fragment including MIDAS and SyMBS metal ions, the MIDAS metal ion coordinating waters (three in state 1 and two in states 3 and 8), and the protein residues in the binding pocket were selected (Supplementary Figure S13). The SyMBS and its coordinating partners were included in the fragment due to the bidentate carbonyl of Glu220 that coordinates both MIDAS and SyMBS. The protein residues in the fragment are truncated using hydrogen capping or methyl/ethyl groups. Using Jaguar,^46,47^ quantum mechanics (QM) calculations were performed to obtain the partial charges for the fragment, which were then fitted and propagated to the receptor to be used in downstream calculations. Restraints were used in fitting the QM-based charges (HF/6-31G* ESP).^39^ Even though the full complexity of electrostatic interactions cannot be captured using static charges, the comparison showed the use of metal custom charge parameterization, despite the absence of the ligand in the QM calculations, improved the classification performance. For the rest of the system, where metal custom charges were not calculated, standard OPLS4 force field parameters were used. Force Field Builder was employed to generate custom parameters for ligand torsions that were not included in the default OPLS4 force field. Unlike the two or three waters at the MIDAS with ZOBs, all other water molecules in AB-FEP simulations were modeled explicitly using the Simple Point-Charge water model.

### Docking

We docked ligands to the closed state (S1) and the open state (S8) using Glide.^48^ For S1, we used the structure prepared from PDB 7TMZ-1 and for S8, we used the structure prepared from PDB 3ZE2-2 (as described above). First, we removed all water molecules that are not connected to ions via ZOBs and generated grids for both open and closed states, centered on the binding pocket. We prepared the ligands using Epik, generating all possible states at pH 7.0 ± 2.0. Then we docked them using Glide in SP (Standard Precision) mode, both the small molecules and the peptides. For each ligand, the top five poses were retained. These were also used to start the subsequent MM-GBSA and — with the exception of linear peptides — the FEP calculations. For the docking-based prediction of whether a ligand is opening or closing, we used the difference of the ligand’s best (lowest) docking score to the open state and the best docking score to the closed state.

### MM-GBSA

MM-GBSA was performed on all the poses generated from Glide (see above) — including peptides — using the VSGB solvation model.^49^ No flexible residues were considered in the calculations. For the MM-GBSA–based prediction of whether a ligand is opening or closing, we used the difference of the ligand’s best (lowest) binding free energy ΔG_open_ to the open state and the best binding free energy ΔG_closed_ to the closed state.

### AB-FEP

We performed AB-FEP calculations for all investigated ligands. When evaluating the performance of our computational models, we used the docked poses generated by Glide. This approach reflects a realistic scenario in which the binding poses of the probed ligands are not known a priori. Only for the two linear peptides, we used experimental poses because the docked poses led to failures in the MD simulations. All AB-FEP simulations were run using FEP+ in the Schrödinger Suite 2024-4.^37,50^ For small molecules, we ran an initial unbiased MD simulation of 1 ns and then an FEP calculation of 10 ns. For the linear peptides, we doubled the initial MD stage to 2 ns and the FEP stage to 20 ns. To keep the protein in its designated conformational state, harmonic position restraints with a force constant of 10 kcal mol^−1^ Å^−1^ were applied to all protein backbone atoms throughout the simulation. In contrast to similar studies on GPCRs,^36^ no flat bottom was used here because of the only very small geometric difference between the states within the ligand-binding pocket. For the AB-FEP–based prediction of whether a ligand is opening or closing, the difference of the ligand’s best (lowest) binding free energy ΔG_open_ to the open state and the best binding free energy ΔG_closed_ to the closed state — analogous to the MM-GBSA–based prediction – were used. We omitted the results from any ligand pose that exhibited bad convergence when other poses of the same ligand converged quickly.

We ran additional simulations to probe the intermediate state S3 as well as the influence of partial charges, ZOBs, and restraints. For these simulations, we only used poses from experimental structures (even for small molecules) which provided the advantage of not having to simulate multiple candidate poses. The only ligands that did not have any structure available were the two M-tirofiban enantiomers. And while it would be less realistic for performance evaluation, this approach is sufficient for gaining methodological insights and for testing hypotheses. To obtain poses for both states, we aligned the experimental structures to the receptor in the probed state and copied over the ligand. As a reference, we repeated the standard workflow on the experimental poses and then ran simulations with the following modifications:

- *State S3*: In addition to S1 and S8, we also probed S3 for which we used the structure prepared from PDB 7UDG-1.
- *Standard charges*: We ran AB-FEP on S1 and S8 using the standard OPLS4 force field without custom modifications to partial charges of the protein, ions, and water molecules.^38–40^
- *Deleted water ZOBs*: All ZOBs connecting ions to water molecules were removed, allowing the water molecules to move freely while the position of the ion relative to the protein was still restrained. Because the force field was parameterized assuming the presence of ZOBs, this approach cannot generally be considered good practice and should be regarded purely as an ad-hoc test.
- *Deleted all ZOBs*: All ZOBs were removed, including those between ions and protein. This means that also the ions moved freely.
- *Flat-bottom harmonic restraints*: We used flat-bottom harmonic restraints instead of the harmonic restraints specified above. Within a range of 1 Å from the restraint center, no force is applied at all. Beyond this range, a regular harmonic potential starts with the same force constant as in the regular protocol (10 kcal mol^−1^ Å^−1^).

### Data Analysis

Each of the tested classification methods provided a classification score for each ligand in the form of a difference of predicted binding free energies or docking scores. While, theoretically, the threshold to classify ligands as opening or closing would be at zero, all of these scores are subject to systematic deviations that require an empirical determination of the threshold.^36^ We thus assessed the predictive power of each classification score using threshold-independent metrics: the areas under the curve of the receiver operating characteristic (AUC-ROC) and of the precision-recall curve (AUC-PRC). Both metrics are calculated using their implementation in scikit-learn.^51^ Plots are generated using matplotlib.^52^ We analyzed molecular trajectories in Maestro and rendered figures in PyMol.^53,54^

## ASSOCIATED CONTENT

### Supporting Information

The Supporting Information is available free of charge on the ACS Publications website.

Supporting figures with more details on methods and analysis and tables with results for all ligands. (PDF)

### Data Availability

The ligand poses, reference files for restraints, numerical results, and analysis scripts are available on GitHub at: https://github.com/schrodinger/integrin-conformational-selectivity

## AUTHOR INFORMATION

### Author Contributions

The manuscript was written through contributions of all authors. All authors have given approval to the final version of the manu-script.

## ACKNOWLEDGMENT

We thank Benjamin Rudshteyn and Edward Harder from the Schrödinger Force Field team and Jarrett Johnson from the PyMol team.

## ABBREVIATIONS

AB-FEP: Absolute Binding Free Energy Perturbation
ADMIDAS: Adjacent to the Metal-Ion Dependent Adhesion Site
AUC-PRC: Area Under the Precision-Recall Curve
AUC-ROC: Area Under the Receiver Operating Characteristic curve
GPCR: G Protein-Coupled Receptor
LFA-1: Leukocyte Function-Associated antigen-
MD: Molecular Dynamics
MIDAS: Metal-Ion Dependent Adhesion Site
MM-GBSA: Molecular Mechanics-Generalized Born Surface Area
PDB: Protein Data Bank
PET: Positron Emission Tomography
QM: Quantum Mechanics
RMSD: Root Mean Square Deviation
SyMBS: Synergistic Metal Binding Site
WT: Wild-Type
ZOB: Zero-Order Bond

