## Supplementary Information for "Conformational Preference Classification of Integrin-Binding Ligands Using Free Energy Perturbation"

### Supplementary Figures

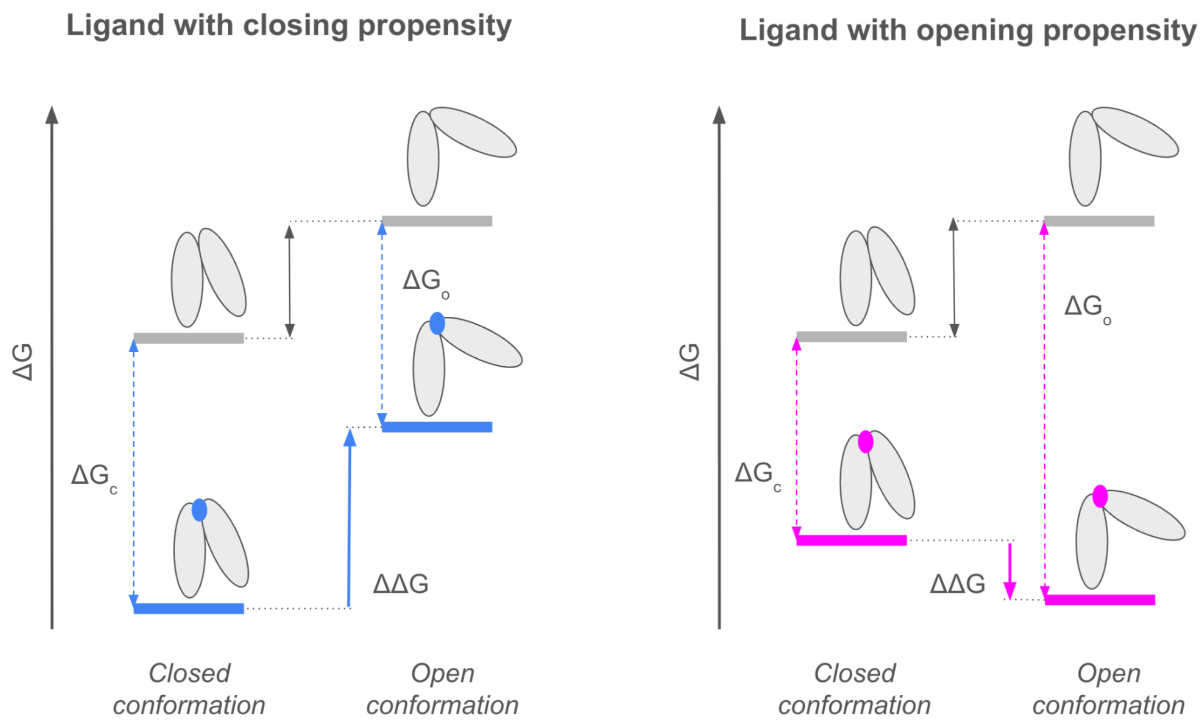


**Figure S1: Translation of closing and opening propensities into binding affinities.** In the apo state (top), the integrin prefers the closed state. Ligands with closing propensity (bottom left) prefer binding to the closed conformation and thus lower the free energy of the closed state more than they lower the free energy of the open state. Ligands with opening propensity prefer binding to the open conformation and thus lower the free energy of the open state more than they lower the free energy of the closed state.

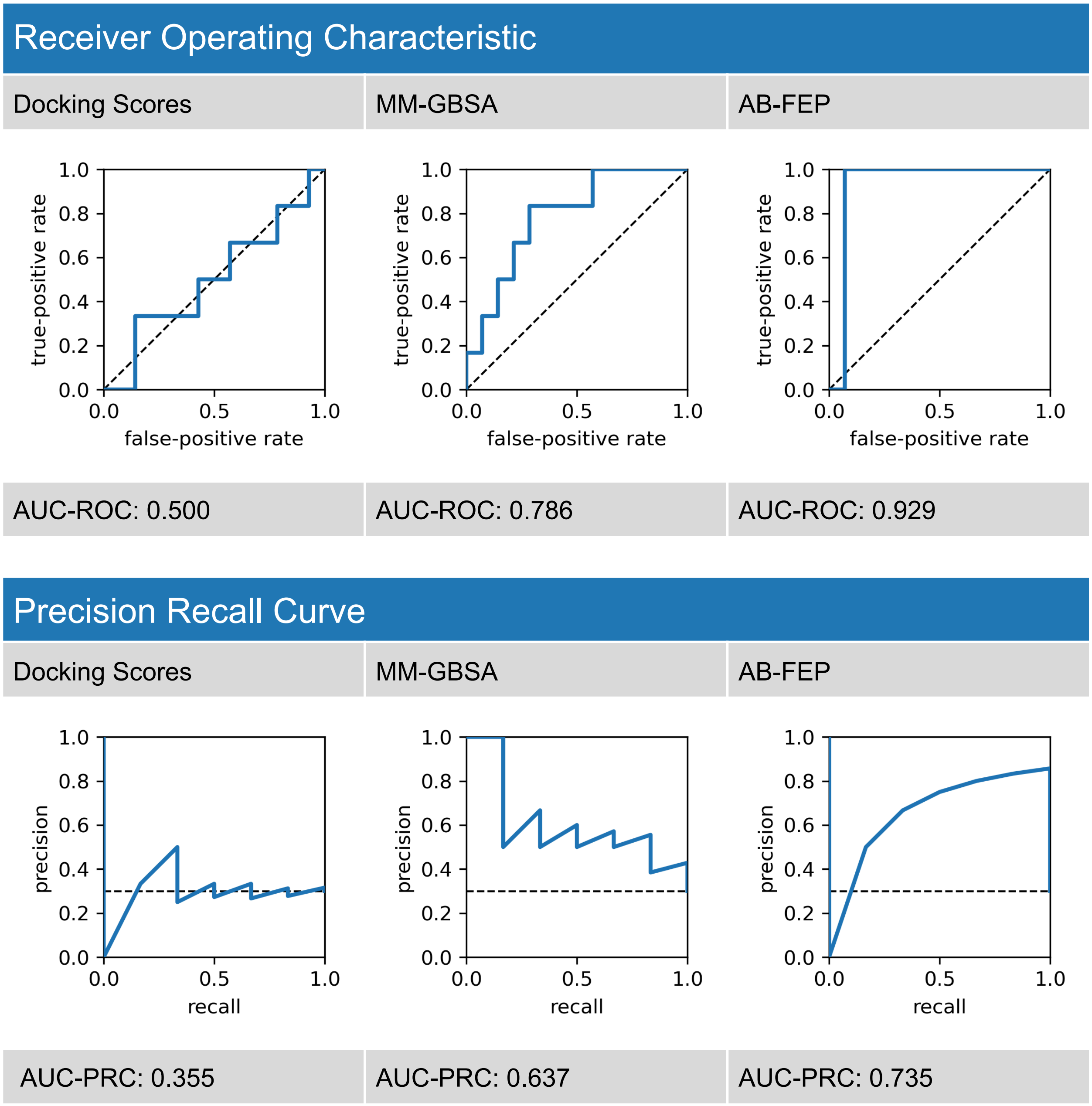


**Figure S2: Classification curves in the performance study** comparing docking scores, free energy estimates from MM-GBSA, and free energy estimates from AB-FEP. From each receiver operating characteristic (ROC) and precision recall curve (PCR), we obtained the corresponding area under the curve (AUC).

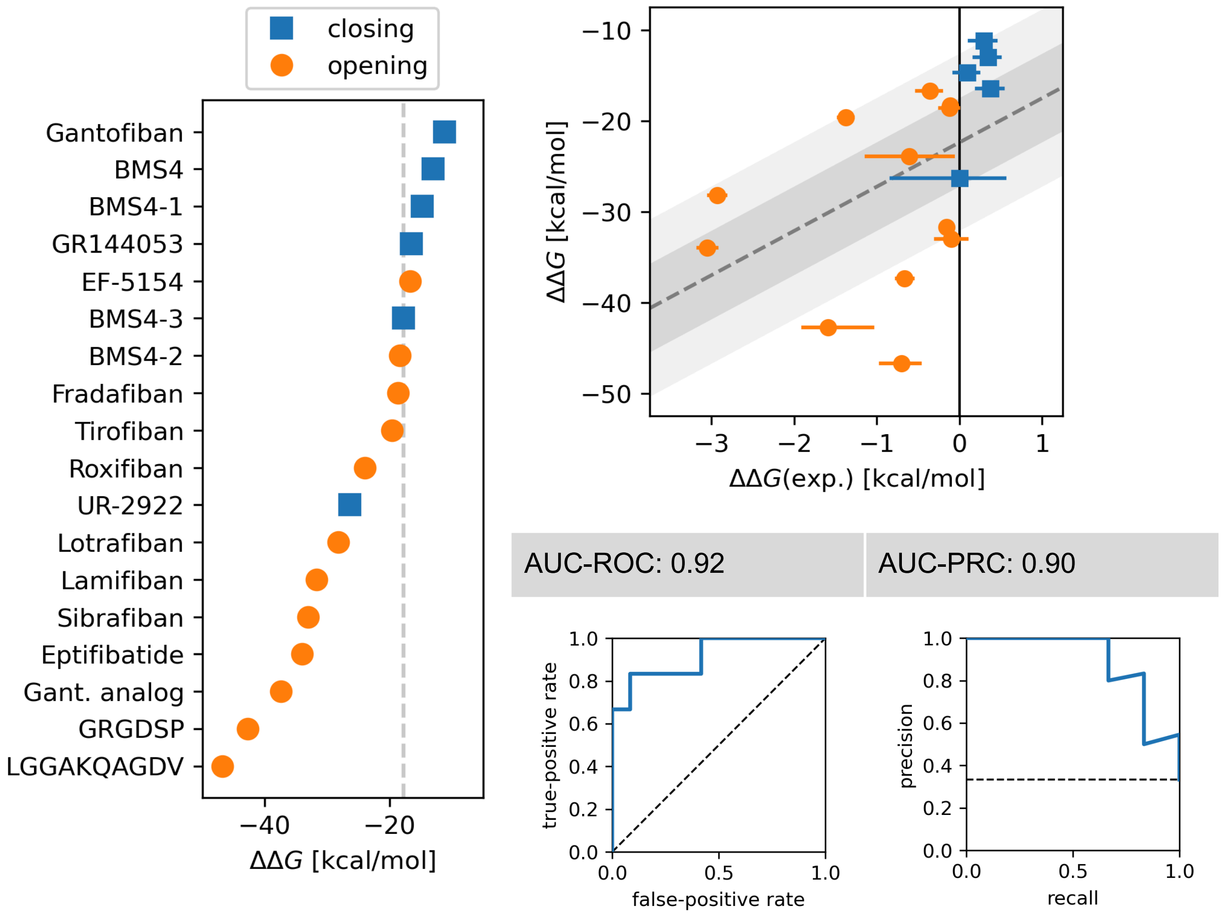


**Figure S3: Results for the AB-FEP workflow on experimental poses.** Left: Free energy difference ΔΔG of binding free energies to the active and inactive state with ligands ordered by the value of ΔΔG. Upper right: Correlation plot between experimental and computational ΔΔG. Lower right: Receiver operating characteristic (AUC) and precision recall curve (PRC), each with the corresponding area under the curve (AUC).

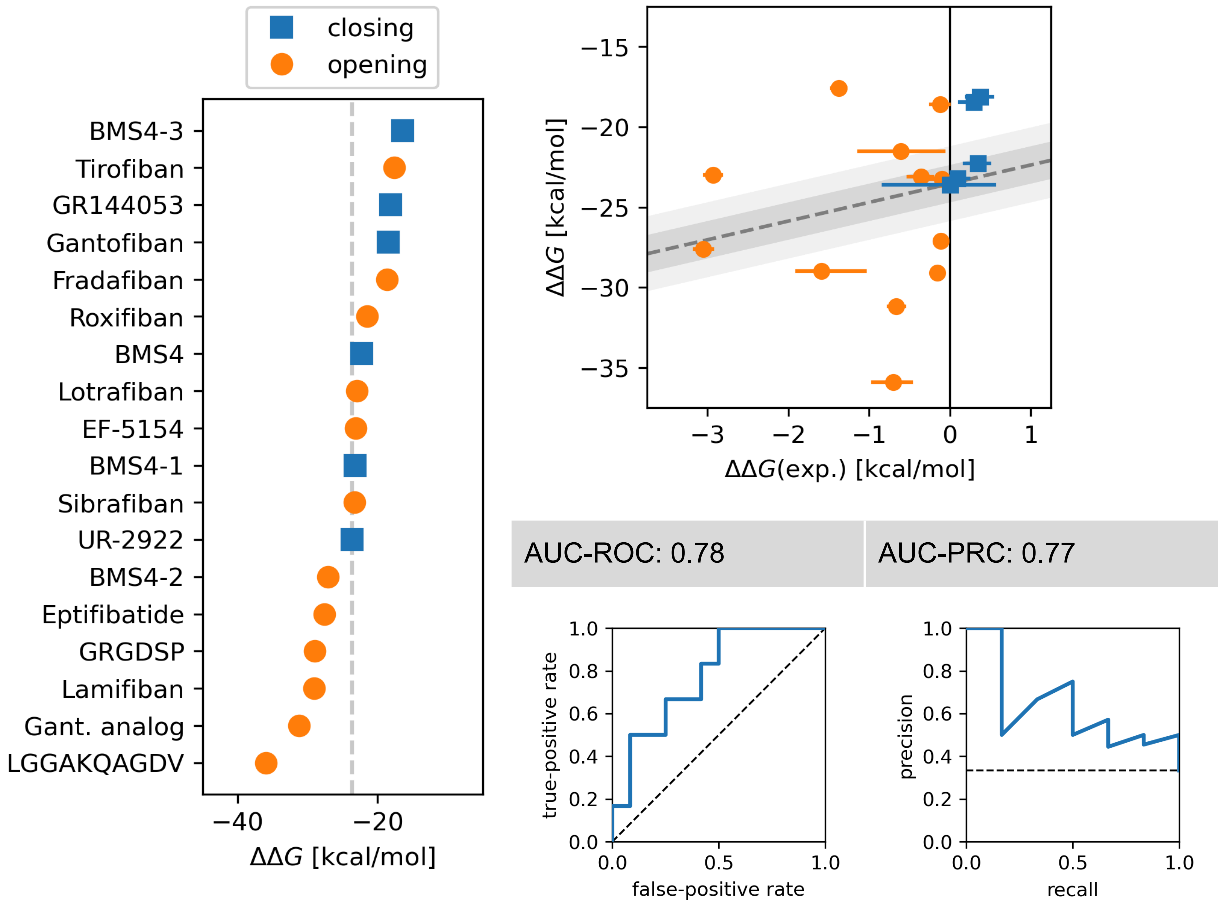


**Figure S4: Results for the AB-FEP workflow with wider backbone restraints.** The half-width of the flat-bottom harmonic restraints was 1 Å. All simulations here were run with experimental ligand poses. Left: Free energy difference ΔΔG of binding free energies to the active and inactive state with ligands ordered by the value of ΔΔG. Upper right: Correlation plot between experimental and computational ΔΔG. Lower right: Receiver operating characteristic (AUC) and precision recall curve (PRC), each with the corresponding area under the curve (AUC).

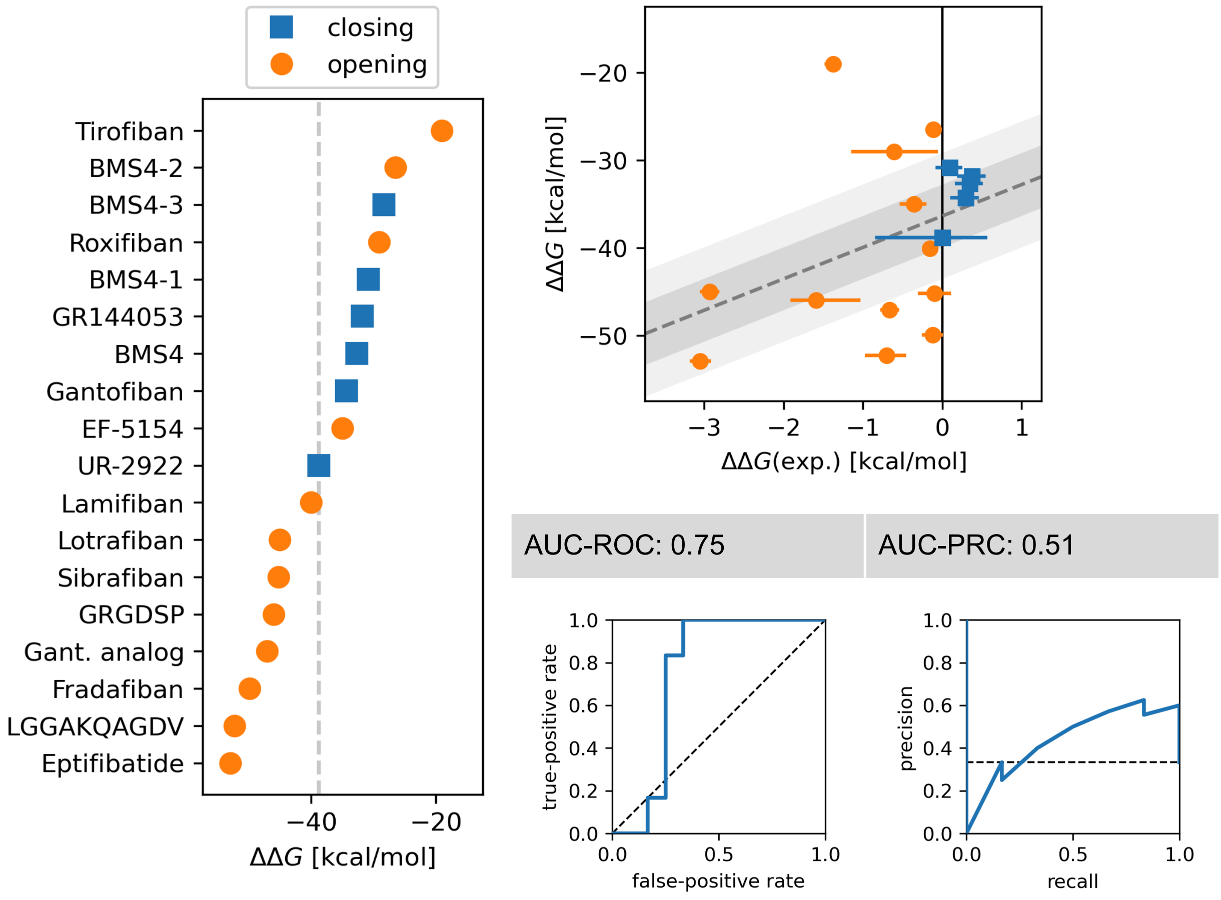


**Figure S5: Results for the AB-FEP workflow with standard OPLS4 charges.** These simulations were performed on PDB 2VDM and 2VDN as the open state S8. All simulations here were run with experimental ligand poses. Left: Free energy difference ΔΔG of binding free energies to the active and inactive state with ligands ordered by the value of ΔΔG. Upper right: Correlation plot between experimental and computational ΔΔG. Lower right: Receiver operating characteristic (AUC) and precision recall curve (PRC), each with the corresponding area under the curve (AUC).

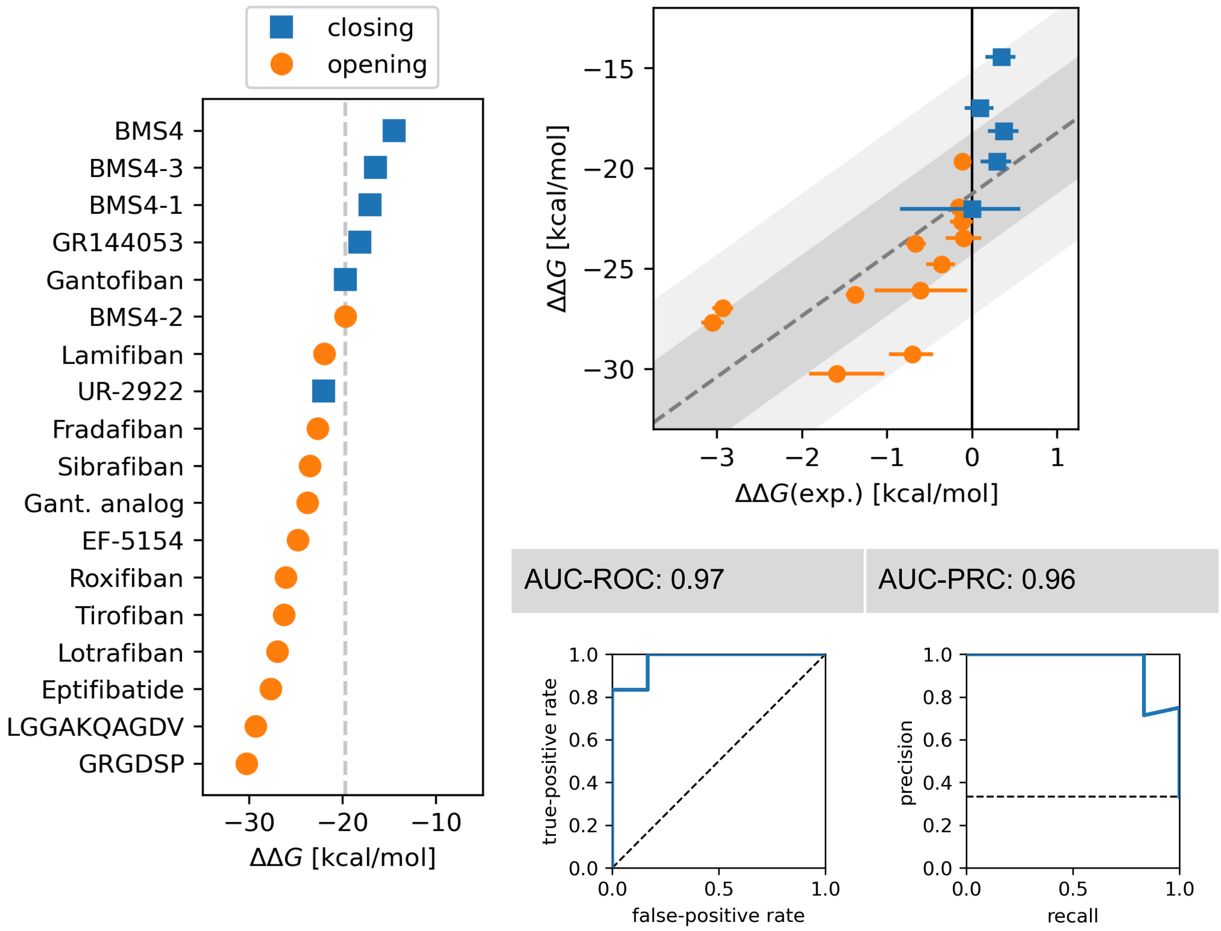


**Figure S6: Results for the AB-FEP workflow without zero-order bonds (ZOBs) to water molecules.** Left: Free energy difference ΔΔG of binding free energies to the active and inactive state with ligands ordered by the value of ΔΔG. Upper right: Correlation plot between experimental and computational ΔΔG. Lower right: Receiver operating characteristic (AUC) and precision recall curve (PRC), each with the corresponding area under the curve (AUC).

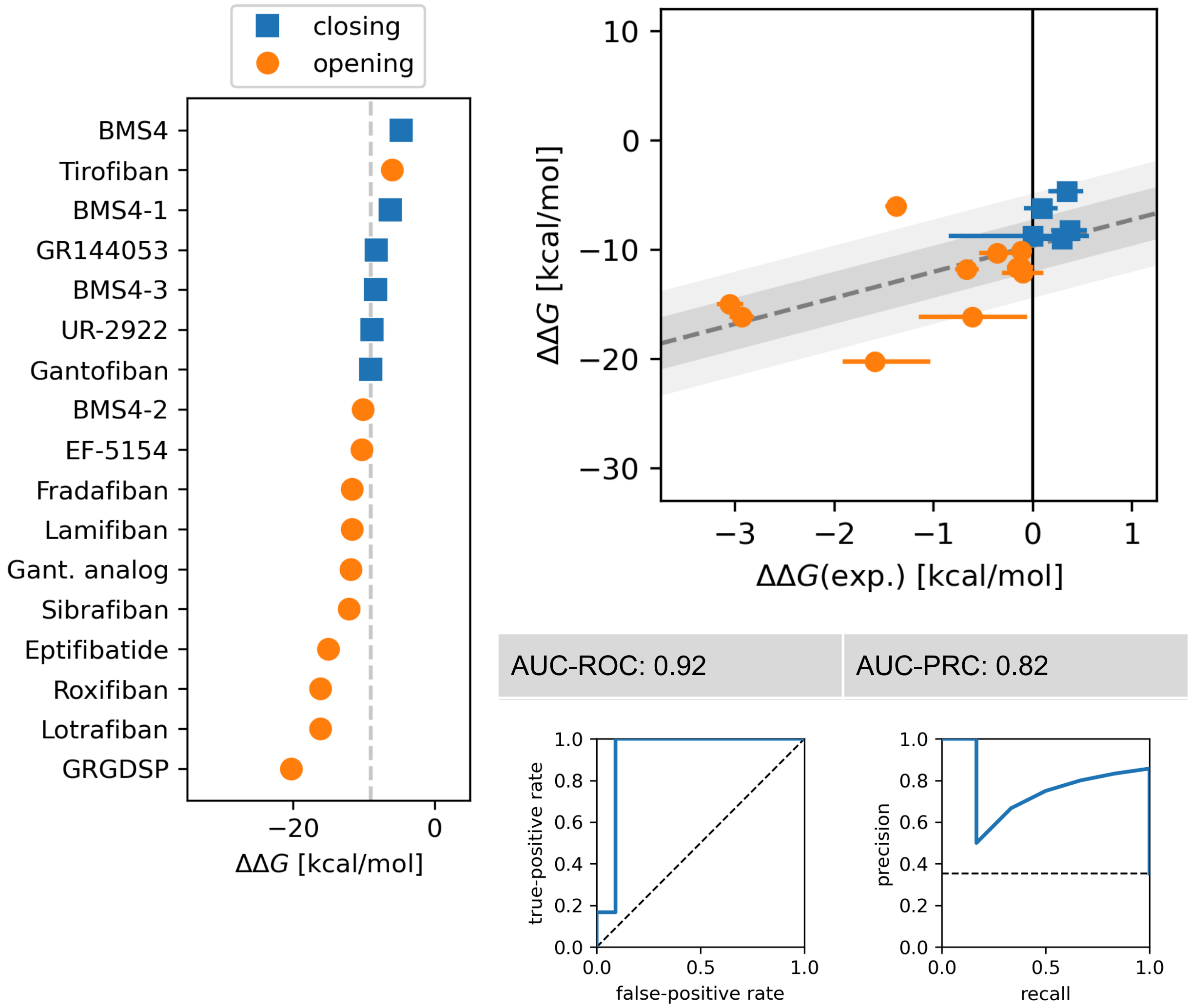


**Figure S7: Results for the AB-FEP workflow without any ZOBs at all.** The open-state simulation failed for the decapeptide LGGAKQAGDV. Left: Free energy difference ΔΔG of binding free energies to the active and inactive state with ligands ordered by the value of ΔΔG. Upper right: Correlation plot between experimental and computational ΔΔG. Lower right: Receiver operating characteristic (AUC) and precision recall curve (PRC), each with the corresponding area under the curve (AUC).

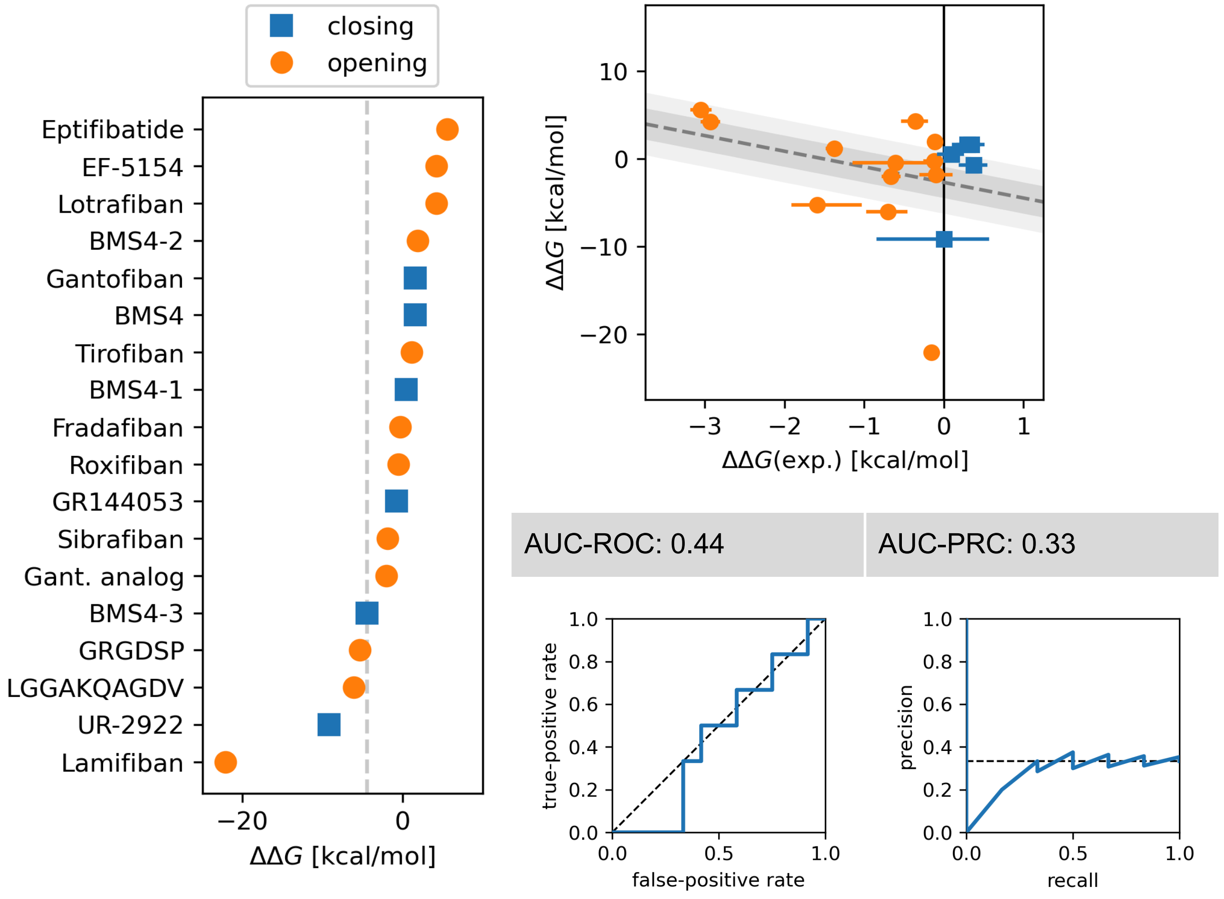


**Figure S8: Results for the AB-FEP workflow using the intermediate state S3 as the closed state.** All simulations here were run with experimental ligand poses. Left: Free energy difference ΔΔG of binding free energies to the active and inactive state with ligands ordered by the value of ΔΔG. Upper right: Correlation plot between experimental and computational ΔΔG. Lower right: Receiver operating characteristic (AUC) and precision recall curve (PRC), each with the corresponding area under the curve (AUC).

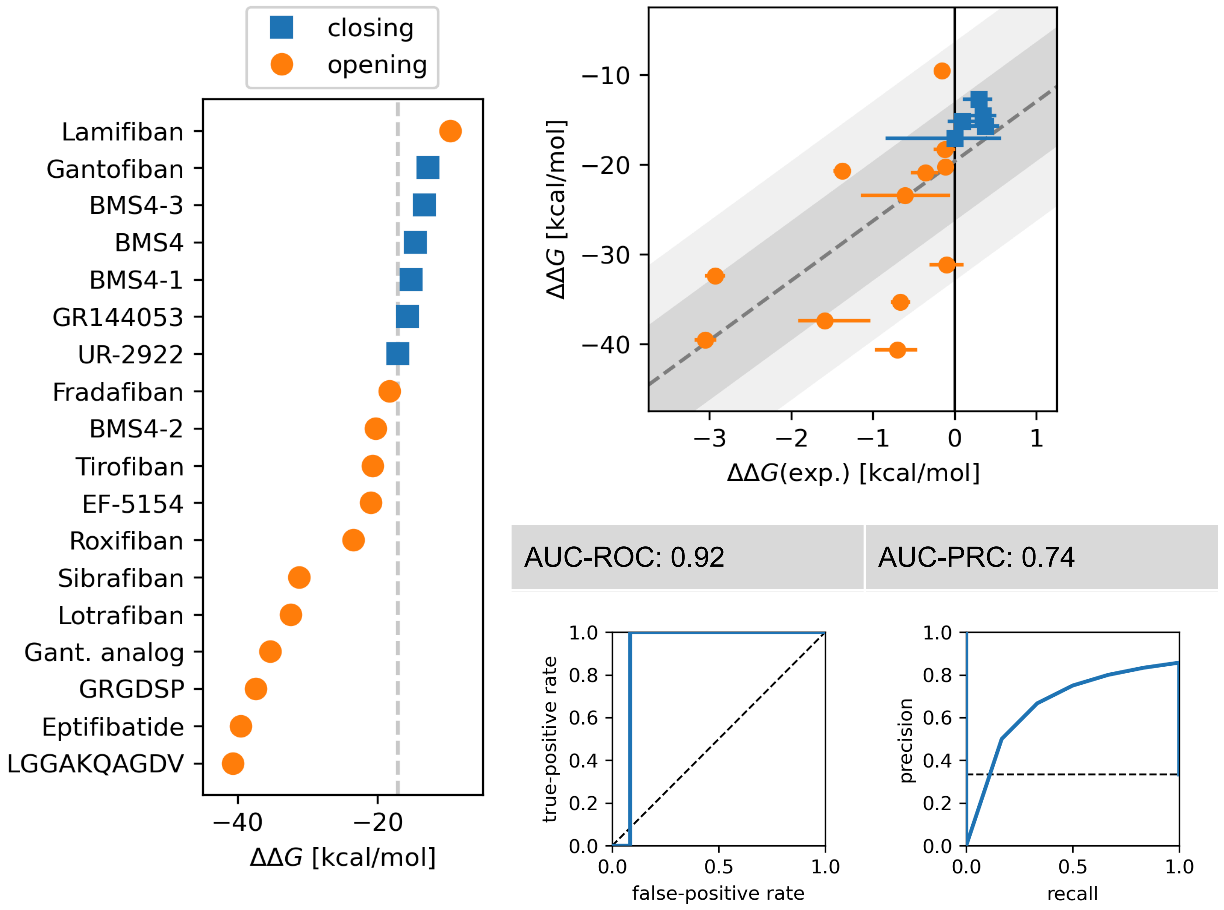


**Figure S9: Results for the AB-FEP workflow using the intermediate state S3 as the open state.** All simulations here were run with experimental ligand poses. Left: Free energy difference ΔΔG of binding free energies to the active and inactive state with ligands ordered by the value of ΔΔG. Upper right: Correlation plot between experimental and computational ΔΔG. Lower right: Receiver operating characteristic (AUC) and precision recall curve (PRC), each with the corresponding area under the curve (AUC).

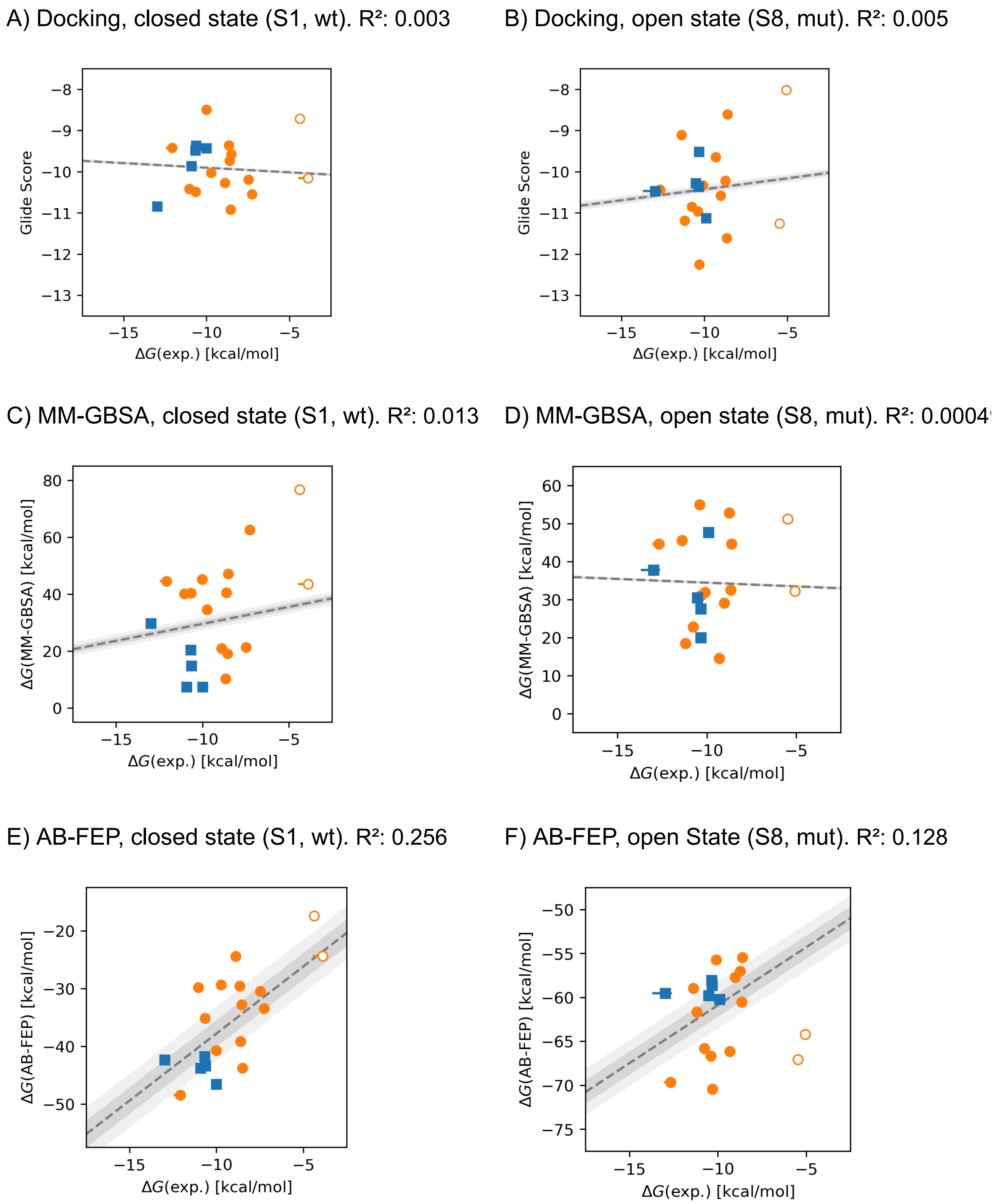


**Figure S10: Affinity scores to individual states S1 and S8 for all three tested methods** (Glide score for docking and binding free energy ΔG for MM-GBSA and AB-FEP) in the performance comparison study and their correlation with experimental values.

Standard AB-FEP Protocol (OPLS4-mcc), started from experimental poses

A) Closed state (S1, wt). $R^2$: 0.16

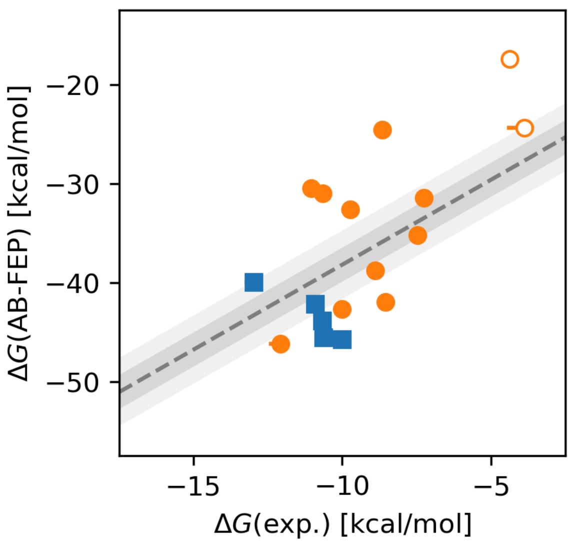


B) Open State (S8, mut). $R^2$: 0.31

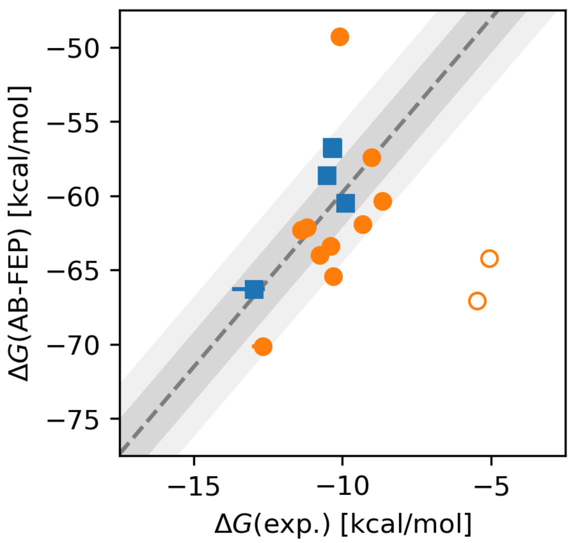


AB-FEP without water ZOBs, started from experimental poses

C) Closed state (S1, wt). $R^2$: 0.51

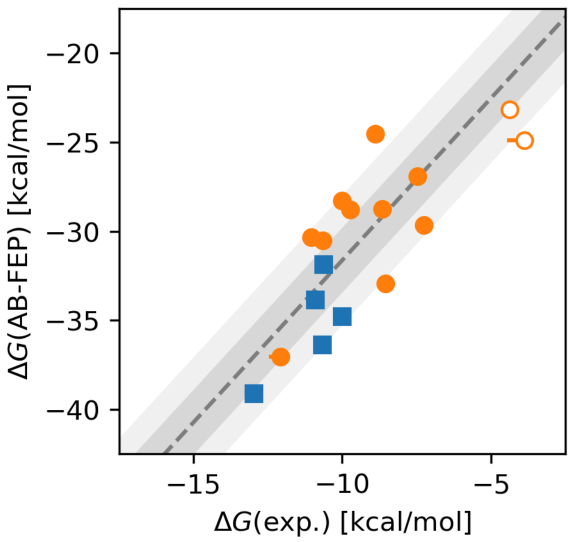


D) Open State (S8, mut). $R^2$: 0.63

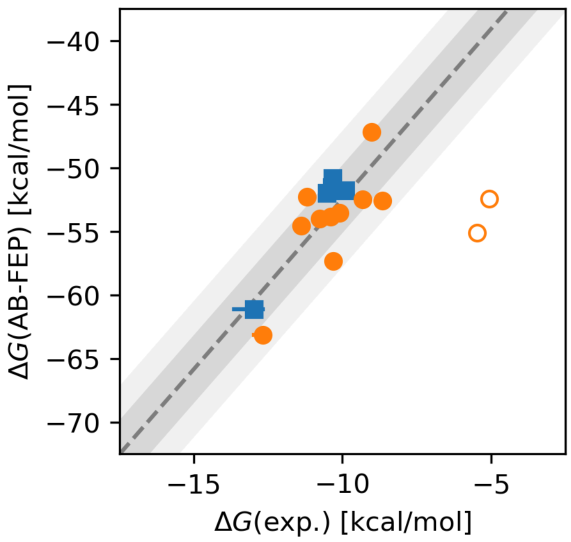


**Figure S11: Binding free energies from AB-FEP to individual states S1 and S8 with and without ZOBs** and their correlation with experimental values. Linear peptides are excluded from the fit and shown with hollow markers.

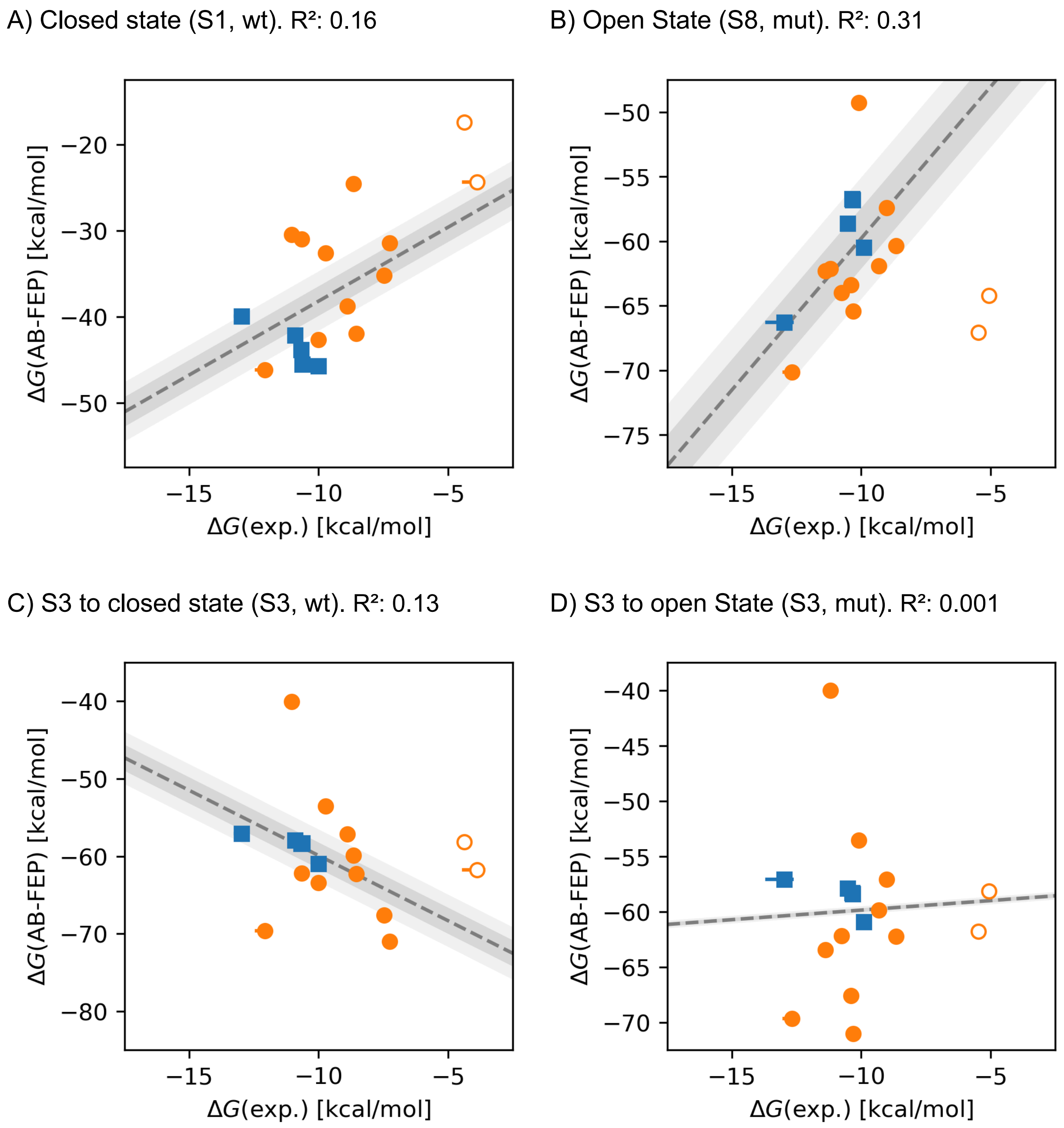


**Figure S12: Correlations of binding free energies from AB-FEP to states S1, S3, and S8** to the experimental results from the wild-type integrin and the mutant (which is stabilized in an open state). These simulations were run using the standard AB-FEP Protocol (OPLS4-mcc) with experimental ligand poses.

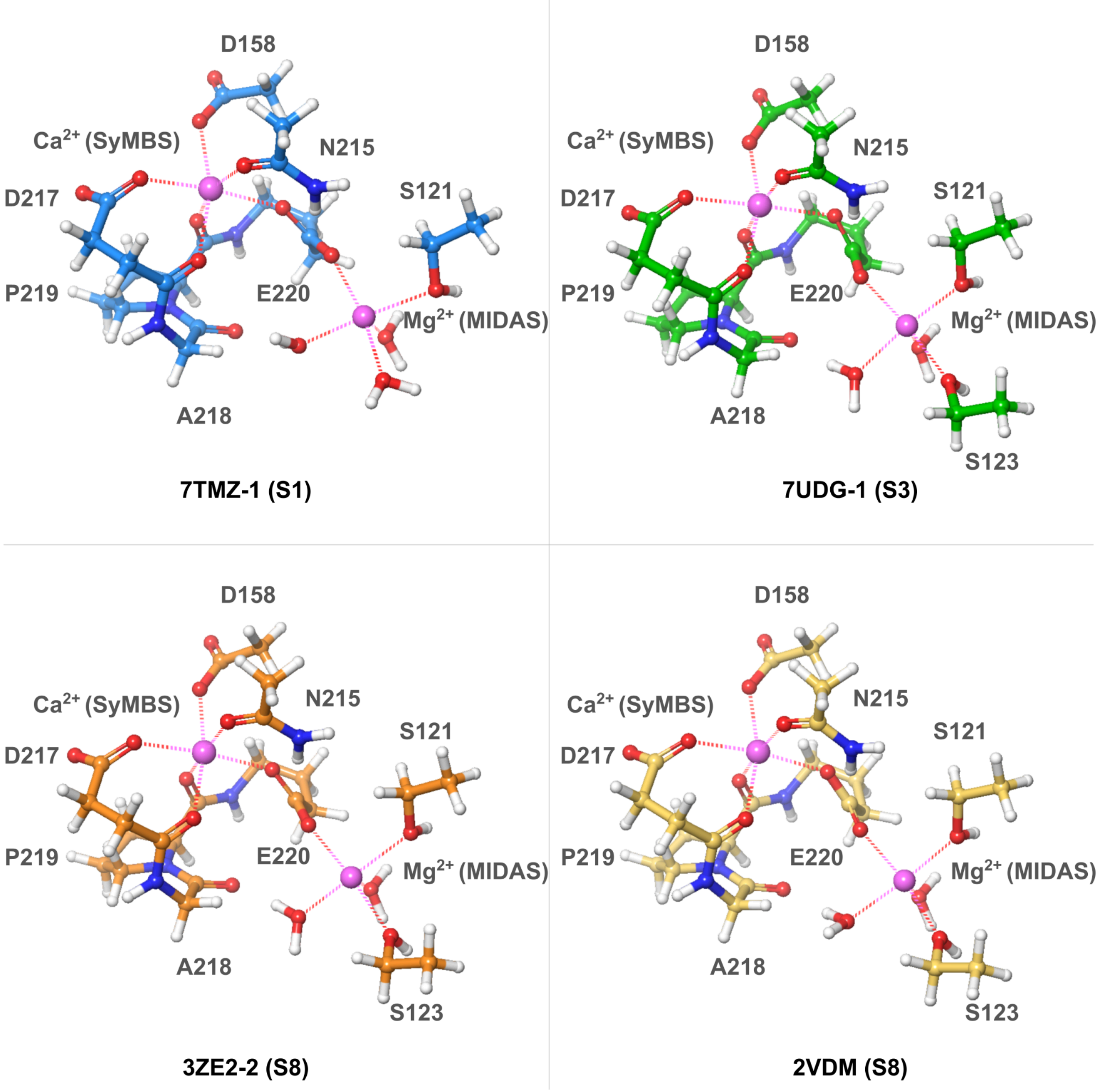


**Figure S13: Capped Fragments** for force field metal custom charge calculations.

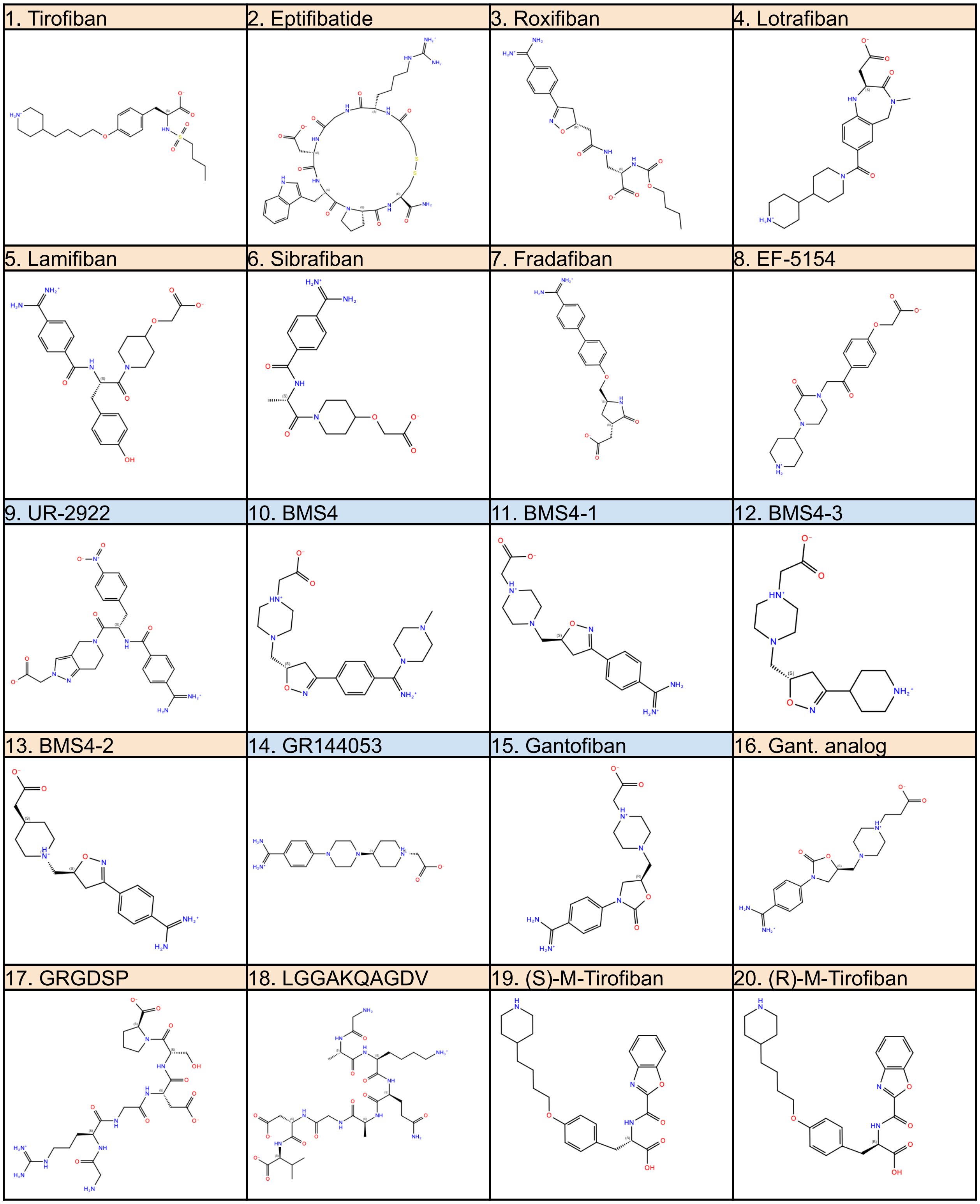


**Figure S14: Chemical structures of the investigated ligands in this study.** Opening ligands' names are shaded in orange and closing ligands' names in blue.

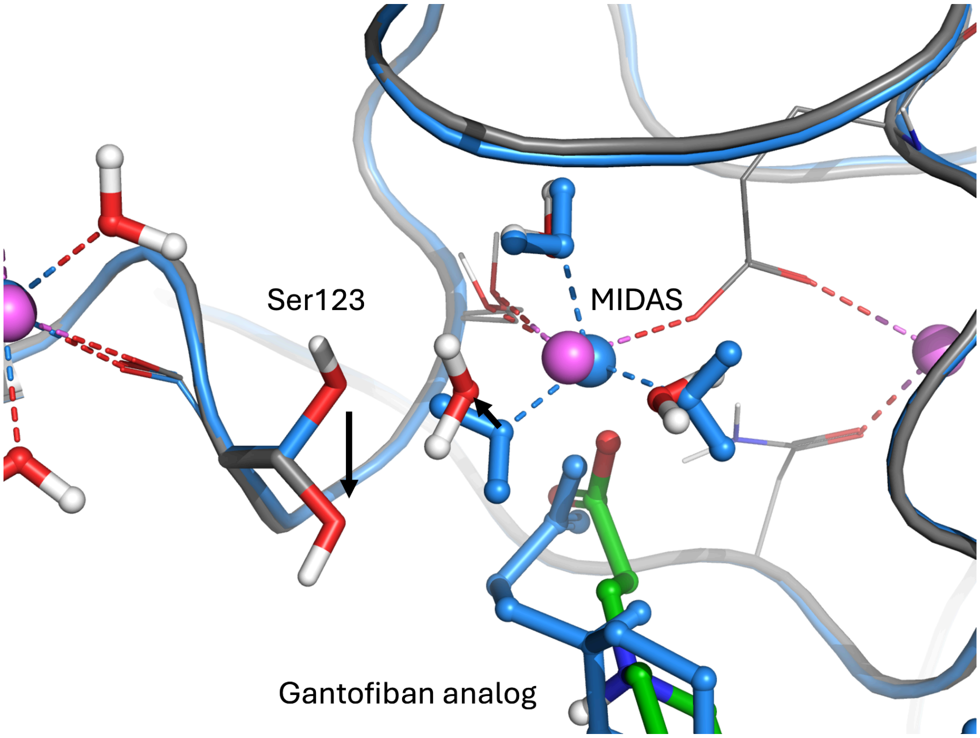


**Figure S15: MIDAS Region without Water ZOBs.** Water molecules around the MIDAS ion in the S1 starting conformation (blue, PDB 7TMZ) and in a representative structure from a simulation without ZOBs (other colors). ZOBs restrain the geometry around the ion and prevent the hydrogen bond network around the ions from exploring alternative configurations like this one that some ligands (here: gantofiban analog) can induce in absence of ZOBs.

A) Experiment (PDB 7TCT) B) Docked to S1 (best ΔG) C) Docked to S8 (best ΔG)

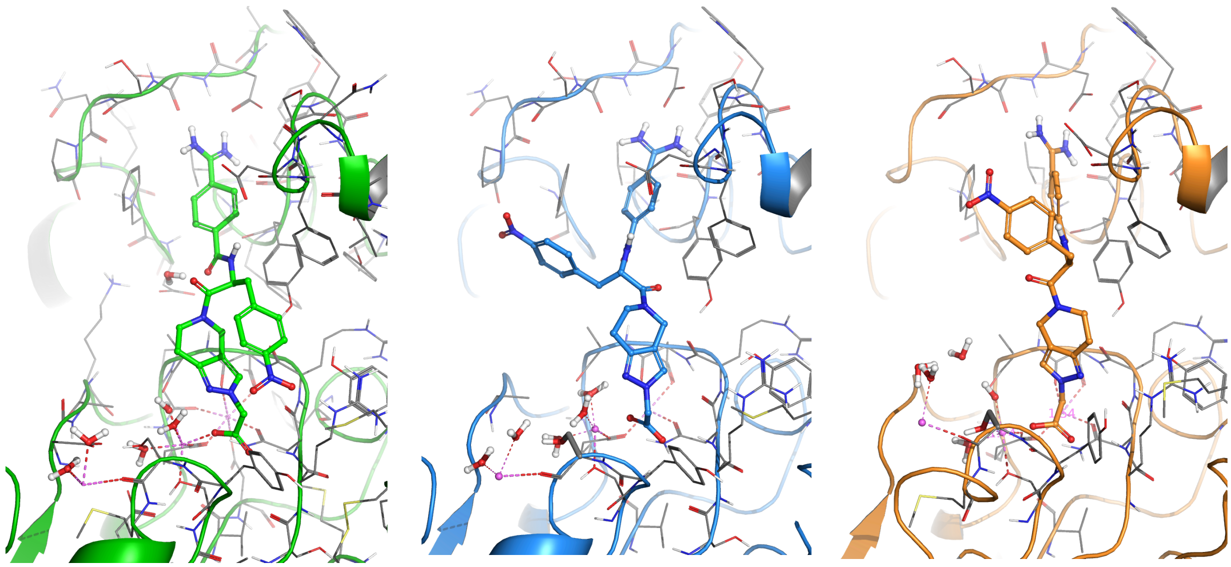

**Figure S16: Poses of UR-2922** from (A) the experimental structure (PDB 7TCT), (B) the docking pose obtained via Glide on state S1 that performed best in AB-FEP, and (C) the best-performing docking pose obtained via Glide on state S8.

### Supplementary Tables

| # | name | opening/ closing | $K_d$ (nM) for αIIbβ3… | | $K_d$(WT)/ $K_d$(N305T) |
|---|---|---|---|---|---|
| | | | WT | N305T | |
| 1 | Tirofiban | opening | 45.9±2.6 | 4.5±0.4 | 10.2±1.1 |
| 2 | Eptifibatide | opening | 4800±500 | 27.7±2.3 | 173±23 |
| 3 | Roxifiban | opening | 1.4±0.6 | 0.5±0.2 | 2.8±1.6 |
| 4 | Lotrafiban | opening | 3300±300 | 23.4±1.9 | 141±17 |
| 5 | Lamifiban | opening | 7.9±0.3 | 6.1±0.3 | 1.3±0.1 |
| 6 | Sibrafiban | opening | 15.1±2.4 | 12.8±2.0 | 1.2±0.3 |
| 7 | Fradafiban | opening | 296±25 | 240±24 | 1.2±0.2 |
| 8 | EF-5154 | opening | 73.2±5.0 | 39.9±7.0 | 1.8±0.3 |
| 9 | UR-2922 | closing | 0.3±0.1 | 0.3±0.2 | 1.0±0.7 |
| 10 | BMS4 | closing | 14.7±1.0 | 26.2±4.9 | 0.6±0.1 |
| 11 | BMS4-1 | closing | 46.2±3.0 | 54±10 | 0.9±0.2 |
| 12 | BMS4-3 | closing | Too low to measure | Too low to measure | – |
| 13 | BMS4-2 | opening | 535±12 | 442±20 | 1.2±0.1 |
| 14 | GR144053 | closing | 10.0±0.8 | 18.8±3.5 | 0.5±0.1 |
| 15 | Gantofiban | closing | 15.8±1.1 | 25.9±5.0 | 0.6±0.1 |
| 16 | Gant. analog | opening | 453±39 | 148±11 | 3.1±0.3 |
| 17 | GRGDSP | opening | $(1.4±0.8)×10^6$ | $(9.6±0.6)×10^4$ | 14.6±8.4 |
| 18 | LGGAKQAGDV | opening | $(6.2±0.8)×10^5$ | $(1.9±0.5)×10^5$ | 3.3±1.0 |
| 19 | (S)-M-Tirofiban | opening | 475±25 | 382±26 | 1.2±0.1 |
| 20 | (R)-M-Tirofiban | opening | 571±19 | 483±22 | 1.2±0.1 |
| – | RUC-4 | neutral | 56.5±3.6 | 55.2±9.9 | 1.02±0.2 |

**Table S1: Ligands considered in this study** with experimental binding affinities to the wild-type (WT) integrin αIIbβ3 and the mutant αIIbβ3 N305T which are used as proxies for the closed and open conformational states, respectively.[31] Ligand 2 is a cyclic peptide (shaded in blue) and ligands 17 and 18 are linear peptides (shaded in purple). All other ligands are small molecules.

| Metric → ↓ State (Struct.) | Dist. [Å] MIDAS –Ser123 | Dist. [Å] MIDAS –Asp126 | Local RMSD [Å] to S1 | Local RMSD [Å] to S8 | Pocket RMSD [Å] to S1 | Pocket RMSD [Å] to S8 | Global RMSD [Å] to S1 | Global RMSD [Å] to S8 |
|---|---|---|---|---|---|---|---|---|
| **S1, closed** (7TMZ-1) | 5.86 | 12.62 | 0.00 | 0.93 | 0.00 | 1.12 | 0.00 | 1.57 |
| **S3, intermed.** (7UDG-1) | 4.41 | 11.58 | 0.74 | 0.46 | 0.51 | 0.83 | 0.21 | 1.53 |
| **S8, open** (3ZE2-2) | 4.18 | 10.63 | 0.93 | 0.00 | 1.12 | 0.00 | 1.57 | 0.00 |

**Table S2: Global vs Local Conformation of 7UDG-1 (S3).** Although the overall shape of the integrin in S3 (PDB 7UDG-1) is closed, the region in the immediate vicinity of the MIDAS ion resembles more the open state. Representative distances of the residue closest to MIDAS (Ser123) show a stronger similarity between S3 and S8. Distances for a slightly more remote residue (Asp126) show a similar difference between S3 and both S1 and S8; here, the intermediate state is right in between open and closed state. RMSD values to the closed and open reference states for the three structures used to represent the open, closed, and intermediate state show strong overall similarity of S3 and S1. The local RMSD was calculated taking into account the residues that are part of the turn of the β6-α7 loop or within a cutoff of 4 Å around the MIDAS ion (Asp119–Ser123, Asn215, Ala218, Glu220, Asp251, Ala252), aligned via the stable β-propeller. The pocket RMSD was calculated taking into account the residues within a cutoff of 5 Å around the ligand, also aligned via the stable β-propeller. The global RMSD was calculated taking into account the β-propeller domain of the α subunit as well as the βI domain and the hybrid domain of the β subunit, aligned via the same components. For reference, the RMSD of the MIDAS ion in a typical AB-FEP simulation is ~0.2 Å and the distance MIDAS–Ser123 fluctuates by ~0.1 Å.

| # | name | opening/ closing | HB network in AB-FEP | |
|---|---|---|---|---|
| | | | Starting from S1 | Starting from S8 |
| 1 | Tirofiban | opening | no | no |
| 2 | Eptifibatide | opening | yes* | yes* |
| 3 | Roxifiban | opening | no | yes* |
| 4 | Lotrafiban | opening | no | no |
| 5 | Lamifiban | opening | no | no |
| 6 | Sibrafiban | opening | no | no |
| 7 | Fradafiban | opening | no | yes* |
| 8 | EF-5154 | opening | no | no |
| 9 | UR-2922 | closing | yes | no |
| 10 | BMS4 | closing | yes | no |
| 11 | BMS4-1 | closing | yes | no |
| 12 | BMS4-3 | closing | yes | no |
| 13 | BMS4-2 | opening | no | no |
| 14 | GR144053 | closing | yes | no |
| 15 | Gantofiban | closing | yes | no |
| 16 | Gant. analog | opening | no | no |
| 17 | GRGDSP | opening | yes* | no |
| 18 | LGGAKQAGDV | opening | yes* | no |
| 19 | (S)-M-Tirofiban | opening | no | no |
| 20 | (R)-M-Tirofiban | opening | no | no |

**Table S3**: **Hydrogen bond network in presence of opening and closing ligands.** Presence of a hydrogen bond network, involving one or two water molecules, bridging the ligand to Ser123, in representative structures of AB-FEP calculations, i.e., the frame that displays the most commonly established interactions during the simulation. Data are shown for AB-FEP starting from docked poses (same as in Figure 5 in the main text) either starting from the S1 or the S8 state. Both sets of AB-FEP simulations show that overall the presence of a hydrogen bond network is a good predictor of whether a ligand is closing or opening. The geometric criteria used to define a hydrogen bond are: H…A distance < 3.00 Å, D-H…A angle > 90° H…A-B angle > 50°. Hydrogen bond networks marked with an asterisk (yes*) are formed with an oxygen atom of the ligand.
